# The past and potential antagonistic effects of fishing ban against nutrient loading control on eutrophication restoration in Lake Hongze

**DOI:** 10.1101/2025.10.05.680530

**Authors:** Bo Qin, Vasilis Dakos, Yongjiu Cai, Yu Zhao, Xiangdong Yang, Enlou Zhang, Rong Wang

## Abstract

Integrated management strategies are often implemented to achieve comprehensive environmental restoration; however, the potential antagonistic effects among such measures are seldom examined. Fishing bans, aimed at enhancing fish biodiversity, may counteract the effectiveness of nutrient control measures in mitigating eutrophication. This study employs the PCLake model and structural equation modeling (SEM) to evaluate this hypothesis in Lake Hongze, the fourth largest shallow lake in China. The PCLake model was calibrated using observational data from Lake Hongze collected between 2016 and 2020. The calibrated model was subsequently applied in a hindcast analysis from 2020 to 2024 and a forecast simulation from 2025 to 2030 to assess the impacts of fishing bans. Combined with SEM path analysis, the results support the hypothesis that fishing bans can exhibit antagonistic effects on nutrient loading control by enhancing top-down regulation relative to bottom-up processes. Specifically, the fishing ban led to increased fish biomass, phytoplankton abundance, and nutrient levels over the past four years, while reducing the abundance of submerged macrophytes, zoobenthos, and zooplankton. Projections further indicate that such antagonistic effects would persist through 2030 if the fishing ban remains in place as planned. Additionally, model forecasts suggest that even with adjustments to the duration and intensity of fishing bans, antagonistic effects could be amplified under more stringent nutrient control scenarios. This study provides valuable insights into maximizing the net benefits of combined environmental management strategies through a systematic framework that evaluates both the effects and underlying processes of different interventions.

## 1. Introduction

Degradations of freshwater lakes, including cultural eutrophication and biodiversity loss, have impeded global sustainable development since the last century (Albert et al., 2020, Ho et al., 2019). Major drivers of these degradations include, but are not limited to, enhanced nutrient enrichment, over-exploitation, hydrological modification, and climate change (Sinha, Michalak and Balaji 2017, Jenny et al., 2020). Achieving relevant Sustainable Development Goals (e.g., Goal 6: Ensure access to water and sanitation for all) requires concerted efforts from multiple stakeholders, implying that integrated management strategies targeting diverse drivers are essential (Birk et al., 2020, Reed 2008). While the synergistic effects of combined management measures have been widely studied, less attention has been given to potential antagonistic effects—which also represent critical information for policy-making.

The shallow lakes in the eastern plain region of China have experienced severe cultural eutrophication and biodiversity degradation (Qin et al., 2013, Wang et al., 2023, Chen et al., 2022). Controlling external nutrient loading is considered the primary approach for mitigating eutrophication, as it restricts phytoplankton growth and influences higher trophic levels through bottom-up resource limitation (Schindler 2012, Jeppesen et al., 2007, Tigli et al., 2024). In China, policies such as the Action Plan for Water Pollution Prevention and Control (2015) and the River and Lake Chief System (2016) are expected to facilitate the restoration of eutrophic lakes as regulations become stricter and more widespread (Yu et al., 2020, Hu et al., 2024). Concurrently, an unprecedented ten-year fishing ban has been implemented since 2020 in the Yangtze River and its connected lakes—one of the most stringent biodiversity conservation initiatives globally (Wang et al., 2022a). This year-round prohibition comprehensively restricts all harvest of natural fishery resources in designated areas. Despite the existence of closed fishing seasons in some lakes—e.g., Lake Hongze from February to June—the permanent withdrawal of all fishing households and revocation of all fishing licenses since 2020 have resulted in a de facto year-round fishing ban in these waters. Classic biomanipulation techniques, such as removing planktivorous and benthivorous fish and introducing piscivorous species, are often applied alongside nutrient control in eutrophic lakes (Søndergaard et al., 2007, Søndergaard et al., 2008). These measures enhance zooplankton biomass by reducing grazing pressure, thereby suppressing phytoplankton proliferation. However, subtropical lakes in eastern China support more omnivorous and planktivorous-benthivorous fish and fewer piscivorous species compared to temperate systems(Jeppesen et al., 2010, Mao et al., 2020). Consequently, the fishing ban may produce effects contrary to traditional biomanipulation: by increasing planktivorous fish stocks, it could intensify predation on zooplankton and indirectly promote phytoplankton growth (Figure 1B). Although recent studies report positive effects in water quality and fish abundance in certain lakes after fishing ban (Huang et al., 2025a, Yu et al., 2020, Zhang et al., 2024), the existence and magnitude of such antagonistic effects remain uncertain—prompting us to ask whether the fishing ban in planktivore-dominated subtropical lakes could paradoxically promote phytoplankton growth by weakening zooplankton grazing pressure, contrary to the outcomes of nutrient control measures.

**Figure 1.**
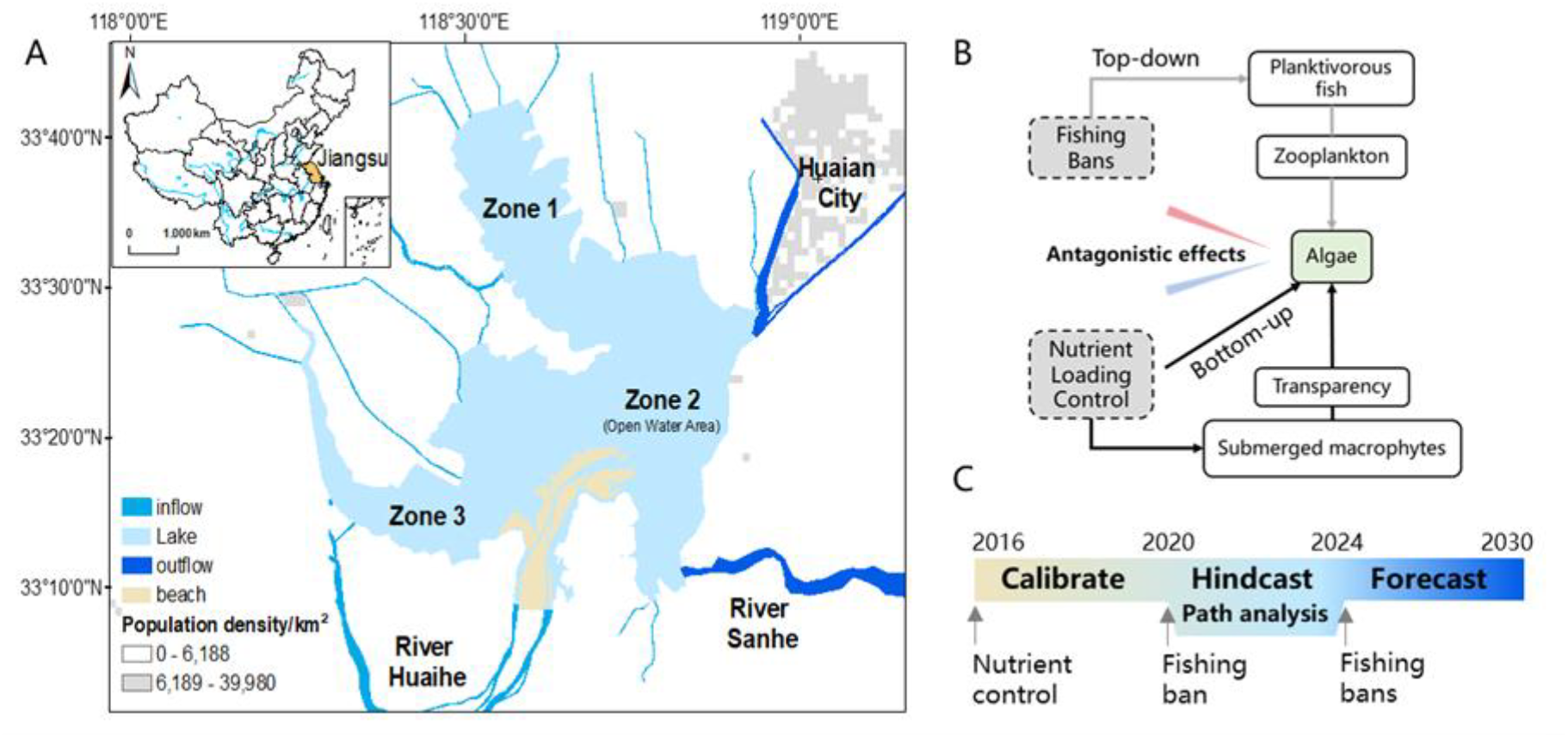
Location of study area, hypothesis and modelling approach of this study.(A) Zones 1 and 3 exhibit relatively high vegetation coverage, whereas Zone 2 is an open water area largely devoid of aquatic vegetation. (B) The conceptual diagram illustrates the hypothesis that fishing bans and external nutrient loading controls may exert antagonistic effects on phytoplankton dynamics, as they operate through distinct ecological mechanisms. (C) Model parameter calibration, hindcast simulation, and forecast analysis will be conducted across different temporal phases. A subsequent structural equation modeling (SEM) path analysis will be performed following the hindcast to elucidate causal pathways.

The process-based lake ecosystem model (PCLake) is an ideal tool to assess such antagonistic effects resulting from fishing bans. The PCLake model has evolved from its original purpose of analyzing eutrophication in temperate shallow lakes into a process-based mechanistic framework for a fully mixed water body. It incorporates key biological components—phytoplankton, zooplankton, and three fish functional groups—alongside nutrient cycling and trophic interactions, with natural feedbacks and producer competition enabling alternative stable states (Janse 2005) (Figure S1). The PCLake model and its derived models (PCDitch, PCLake+, and PCLakeS+) has been widely applied across diverse geographic regions and climatic zones to address a broad range of ecological and management questions (Janssen et al., 2015). Key application areas include identifying critical nutrient loading thresholds for lake degradation and restoration (Qin et al., 2022,Wu et al., 2024), assessing lake resilience under climate warming (Qin et al., 2025, Rolighed et al., 2016), exploring the role of hydrological variability in driving regime shifts (Kong et al., 2017, Xue et al., 2025), and developing early-warning indicators for impending regime shifts (Gsell et al., 2023). Furthermore, the model has been employed to examine food web stability and the interactions among functional groups, including fish and other biological components (Zhang et al., 2023). In terms of applicability, the models are suitable for various types of aquatic systems, including fully mixed shallow lakes, ditches (e.g., PCDitch), and stratifying lakes (e.g., PCLakeS+). Model developments have incorporated key ecological drivers such as thermal stratification, spatial heterogeneity, climate variability, and lake–catchment connectivity, thereby expanding their scope to a wide range of environmental conditions and management scenarios.

This study employed two modeling approaches to evaluate the potential antagonistic effects between nutrient control measures and fishing bans in Lake Hongze, the fourth largest shallow lake in China (Figure 1). As a key storage reservoir for the Eastern Route of the South-North Water Diversion Project, Lake Hongze plays a vital role in climate regulation, flood control, and water resource management, and is currently in a critical phase of restoration (Zhang et al., 2024). Monthly observational data from 2016 to 2020, collected prior to the implementation of the fishing ban, were used to calibrate the PCLake model. A hindcast simulation based on the calibrated model was then conducted to assess the ecological impacts of the fishing ban, the trophic cascades of which were further examined using structural equation modeling (SEM). SEM is used here not for causal inference, but to empirically evaluate the strength of associations and to test pathways derived from the mechanistic PCLake simulations. A forecast analysis was performed to identify strategies for maximizing management effectiveness. The results confirm the hypothesized antagonism using a novel research framework, and both the implications and limitations for lake conservation and management are discussed.

## 2. Material and methods

### 2.1 Study Area

Lake Hongze (33°06’-33°40’N, 118°10’-118°52’E), the fourth largest freshwater lake in China, has a surface area of 1793 km^2^ and a drainage basin of 160,000 km^2^. With a mean depth of 1.7 m and a water volume of 3.0 billion m^3^ (Cai et al., 2016a), the lake exhibits a rapid hydrological cycle and a short water retention time of approximately 35 days. The region experiences a monsoon-influenced climate, with a mean annual precipitation of 925.5 mm, an average wind speed of 3.7 m·s^-1^, and mean temperatures of 2.5°C in January and 28.8°C in July. Major inflows enter from the west and exit toward the east (Figure 1A; Table S1). The Huaihe River, the primary inflow, accounts for about 70% of total inflow and more than half of the external nutrient load. The main outflow occurs through the Sanhe River, which connects Lake Hongze to the Yangtze River and conveys 60%–70% of the total discharge. The fish community in Lake Hongze has been dominated by zooplanktivorous, benthosivorous, and omnivorous species, while predatory fish represent less than 25% of the total (Table S2).

Spatial heterogeneity characterizes the lake’s water quality and ecological conditions (Figure 1A, S2). Zone 1 (Chengzi District) in the north has restricted hydrologic exchange, while Zone 3 (Lihewa District) in the west functions as a wetland reserve with shallow water. These regions support relatively abundant populations of aquatic macrophytes, zooplankton, and macroinvertebrates (Li et al., 2019, Ni et al., 2023, Cai et al., 2016b). In contrast, Zone 2 is an open-water area largely devoid of submerged vegetation and exhibits higher nutrient levels, strongly influenced by inputs from the Huaihe River.

Since the 1980s, Lake Hongze has undergone eutrophication following marked declines in water quality and aquatic biodiversity (Li and Pu 2003). It was listed under the Regulations of Jiangsu Province on Protection of Lakes in 2004 to mitigate pollution and manage resource use. Between 2017 and 2019, nine algal blooms were recorded, with affected areas expanding from 16 km^2^ to 36 km^2^ (Cui et al., 2021).

### 2.2 Sampling and Measurement of Lake Variables

Monthly field investigations were conducted from January 2016 to December 2019 at ten sampling sites distributed across three zones (Figure 1A). These sites are characterized by very shallow conditions, with annual average water depths ranging between 1.9 m and 2.9 m. At each site, Secchi depth (SD) was measured in situ, and dissolved oxygen was measured using a YSI 6600 V2-4 Multi-Parameter Water Quality Sonde. Vertically integrated water samples were collected using acid-cleaned 5 L plastic containers, stored under cool and shaded conditions, and transported to the laboratory for subsequent analysis. Total nitrogen, total phosphorus, and chlorophyll-a concentrations were measured following standard methods described in APHA (2012). Zoobenthos samples were collected at all sites, where the substratum predominantly consisted of muddy sediment. Sampling was performed using a 0.05 m^2^ modified Peterson grab, with three grabs combined to form a composite sample. The collected material was sieved in situ through a 250 μm mesh sieve. Retained residues were stored in a low-temperature incubator and transported to the laboratory on the same day. There, samples were sorted on a white tray, and organisms were preserved in 7% buffered formalin. Specimens were identified to the lowest practicable taxonomic level, enumerated, blotted dry, and weighed for wet mass determination using an electronic balance (Sartorius BS-124, precision: 0.1 mg). Taxonomic identification and classification of zoobenthos followed the references of Liu et al. (1979) and Tang (2006).

### 2.3 Lake Ecosystem Model Configuration and Calibration

To achieve adequate model performance for Lake Hongze during 2016–2024, key model inputs were adjusted. Fixed inputs included a mean water depth of 1.9 m, maximum depth of 4.5 m, wind fetch of 60,600 m (Janssen et al., 2014), lightly clayish soil (Janse 2005) and no wetland zone. Time-series inputs of PCLake model, including wind speed (m·s^-1^), solar radiation (J·m^-2^·d^-1^), water temperature (°C), water inflow, evaporation (mm·d^-1^), outflow (mm·d^-1^), and nutrient loading (g·m^-^ 2·d^-1^), were derived from multiple sources: (1). Monthly wind speed, precipitation, air temperature, and radiation from 2016 to 2024 were obtained from National Meteorological Information Center (https://m.data.cma.cn/data) at a meteorological station (Xuyi, 32.98°N,118.52°E, 40.8 m a.s.l) located 3 km from the lake. (2) Water temperature from the Huaihe River (2016–2019) was used directly; an empirical air–water temperature relationship extended these data to 2020–2024. (3). Monthly river inflow (2016–2019) was investigated (Table S1), with the Huaihe River contributing 88.5% of total inflow. Inflow for 2020–2024 was estimated using precipitation and historical discharge. Daily water level (Jiangba station) ranged from 11.5 m to 13.7 m. Mean daily evaporation was set to 2.5 mm (variance: 1.3 mm). Outflow was computed from total inflow, water level change, and evaporation. (4). Monthly nutrient inputs (2016–2021) from the Huaihe River averaged 4×10^5^ kg N and 2×10^4^ kg P (Meng et al., 2023), showing a general decline, especially after December 2018. A ratio of 0.8 was applied to estimate total nutrient loading based on Huaihe inputs (Ji et al., 2014).

Two modifications were made to the PCLake model to better represent Lake Hongze’s ecosystem and address the research objectives. First, while the original model includes three functional fish groups—zooplanktivorous juvenile whitefish, benthivorous adult whitefish, and piscivorous fish—with benthivorous adults (sDFiAd) dominating (∼80% in both clear and turbid states), we introduced an additional group: zooplanktivorous adult whitefish. This modification was motivated by the observation that zooplanktivorous and omnivorous fish dominate local catches, a pattern that is not simply a result of high juvenile abundance, but rather reflects the presence of adult individuals that remain in the zooplanktivorous niche throughout their life cycle. In the original model, the juvenile stage is only temporarily zooplanktivorous and lacks the longevity and biomass accumulation characteristic of a true adult zooplanktivorous population. Therefore, simply increasing the growth or reproductive rates of juveniles would not adequately represent the persistent top-down control exerted by these adult fish. All equations for this group were adapted from those of benthivorous adults, except for the food limitation function, which was derived from zooplanktivorous juveniles. This design ensures that the original juvenile parameterization—well established in the PCLake literature—is preserved, while the observed fish composition is explicitly represented in the model structure. Second, the original model represents fishing intensity as a first-order rate constant, which is inadequate for simulating variable fishing pressures. To improve temporal realism, we converted fishery intensities into time-series inputs based on annual fish yield and composition data (Mao et al., 2019), allowing dynamic simulation of fishing effort over time.

Default model parameters were initially derived from observations in over 40 lakes (Janse et al., 2010). Both original and newly introduced parameters were adjusted to enhance the performance of the modified PCLake model for Lake Hongze. Calibration was performed over a four-year period (2016–2020) using monthly observed data for total nitrogen, total phosphorus, dissolved oxygen, Secchi depth, chlorophyll-a, and zoobenthos, along with annual records of macrophyte coverage and fish harvest from published sources. Although the vegetation presence frequency (VPF) ranges between 0.24 and 0.37 (Li et al., 2019), submerged macrophytes remain scarce, with a peak coverage of only 11.80 km^2^ (approximately 0.8% of the lake area) between 2013 and 2023 (Huang et al., 2025b). Fish catch data from 2016 to 2019 were provided by the Jiangsu Province Hongze Lake Fisheries Management Committee (https://www.jiangsu.gov.cn/art/2015/9/16/art_46143_2543010.html). The zooplanktivorous and omnivorous fish collectively accounted for over 75% of the total catch (Lin. et al., 2013, Mao et al., 2019).

Simulations were conducted using a C++ compiled version of PCLake called via GRIND for MATLAB (Mooij et al., 2014). Each simulation began from a default turbid state and was run for a 20-year “burn-in” period under 2016 input conditions to achieve seasonal equilibrium before calibration. Key parameters were manually calibrated based on model outputs. The coefficient of determination (R^2^) and root-mean-square error (RMSE) were employed to evaluate model performance across whole-lake averages (Bennett et al., 2013). A higher R^2^ indicates better predictive accuracy, while a lower RMSE reflects smaller deviations between simulated and observed values. Given the absence of abrupt ecological shifts during the calibration period, the objective was to achieve simulations where most variables exhibited similar ranges and seasonal dynamics to the observations.

### 2.4 Hindcast and SEM analysis

Following calibration, the model simulation was extended from 2020 to 2024 to conduct a hindcast analysis. Due to the unavailability of nutrient loading data for this period, values from 2020 were applied to the subsequent four years. Fishing intensity was set to zero to reflect the implementation of the fishing ban (FB scenario). For comparative purposes, a no-fishing-ban scenario (no FB) was also simulated using the average fishing intensity from 2016 to 2020. We excluded dissolved oxygen from the analysis, given its weak regulatory effect on the food web of this large, shallow system. By contrast, zooplankton were included due to their pivotal role as a trophic link between fish and phytoplankton, which is central to the ecosystem dynamics of Lake Hongze. Although additional time-series observations were unavailable, further validation was supported through comparison with previously published findings in the discussion.

Although PCLake is built upon a rigorous theoretical framework, the complexity of biological processes often complicates direct and concise representation of trophic cascades. In contrast, structural equation modeling (SEM), grounded in a strong theoretical foundation, allows for the estimation of trophic cascade strength and other causal networks through a series of regression analyses (Grace 2006, Su et al., 2019). Thus, integrating both approaches enables a comprehensive assessment of both the outcomes and underlying processes associated with combined management strategies. Structural equation modeling (SEM) was employed to analyze trophic cascades under scenarios with and without the fishing ban. A conceptual path model was developed based on the PCLake framework (Figure S1), incorporating seven variables. Both total phosphorus and total nitrogen are simulated in the two nutrient loading scenarios. However, because total nitrogen was closely correlated with total phosphorus, we incorporated only total phosphorus into the conceptual path model. Monthly outputs from both simulations were used to ensure sufficient data. The model was fitted to the covariance structure of the data, beginning with a full model that included all possible pathways, which was subsequently refined by removing nonsignificant paths. Standardized path coefficients were used to quantify the strength and direction of direct relationships between consecutive trophic levels. The overall cascade strength was evaluated through the indirect effect from top to bottom of the food web, derived from the product of sequential path coefficients. Model fit was assessed using the chi-square/degrees of freedom ratio (CMIN/df) (Karin Schermelleh-Engel and Helfried Moosbrugger 2003), to assess how well the proposed network structure explains the simulated data from PCLake. All SEM analyses were performed in AMOS 21.0 (IB12).

### 2.5 Forecast

According to the Environmental Quality Standards for Surface Water in China, the total phosphorus level in Lake Hongze in 2023—though the lowest in recent years—still exceeded the drinking water threshold by 46% (Zhang et al., 2024). In addition, while fish stocks increased significantly, imbalanced fish community structure was also reported. Above facts imply that continued measures are further needed for Lake Hongze in the near future. To evaluate potential management outcomes, we designed eight scenarios combining different levels of external nutrient loading and fishing intensity to simulate conditions from 2025 to 2030. Two nutrient loading scenarios were defined: the highest level over the past five years (NP-, control) and the lowest level (NP--, approximately half of NP-). Four fishing scenarios were established: continued full fishing ban (F0, control); a biennial ban alternating between prohibition and permission (F0/1); full resumption of fishing at the 2020–2024 average intensity (F1); and continuous fishing at half of F1 intensity (F0.5). Other model inputs used 2020 data as boundary conditions. As the ten-year fishing ban is scheduled to end in 2030, we focused on the relative changes between 2030 and 2025. Differences in annual model outputs were calculated and compared against the control scenario to standardize units across variables.

## 3. Results

### 3.1 Lake States and PCLake Model Performance

The average nutrient concentrations in the lake water from 2016 to 2020 were 1.72 g·m-3 for total nitrogen (TN) and 0.08 g·m-3 for total phosphorus (TP) (Figure S2). Total phosphorus (TP) exhibited a slight upward trend, whereas total nitrogen (TN) remained broadly stable. The mean Secchi depth was 0.35 m and dissolved oxygen averaged 0.35 g·m-3, both remaining relatively stable throughout the period. The chlorophyll-a concentration averaged 8 mg·m-3, with occasional peaks observed. At the class level, Oligochaeta dominated in Zone 1, whereas Polychaeta were more abundant in Zone 2 (Figure S3). Since the PCLake model does not explicitly include large zoobenthos, the total dry weight of the five major classes was used to represent zoobenthos biomass.

The performance of the calibrated PCLake model for Lake Hongze was assessed using two statistical indices based on six monthly and two annual observational variables (Figure 2, Figure S4). The simulated trajectories (solid lines) generally captured both the range and seasonal dynamics of the whole-lake observations (dots). The coefficient of determination (R^2^) ranged from 0.07 (zoobenthos) to 0.91 (dissolved oxygen), with an average of 0.46. The root-mean-square error (RMSE) varied between 0 (fish catch) and 3.63 (chlorophyll-a), averaging 0.70. The zero RMSE for fish catch reflects the very small magnitude of the observed data (typically near zero), and rounding to two decimal places in the figures resulted in values of zero. Although the annual observations (macrophyte coverage and fish catch) were limited in number, their simulations were deemed acceptable due to agreement in mean values. Monthly water quality variables were better reproduced than biological variables (phytoplankton and zoobenthos). The modified and calibrated model generally exhibited higher R^2^ and lower RMSE compared to the original model (Figure S5, Table S3), particularly for biological components. Overall, the model demonstrated acceptable performance during the 2016–2020 calibration period and was considered suitable for further analysis.

**Figure 2.**
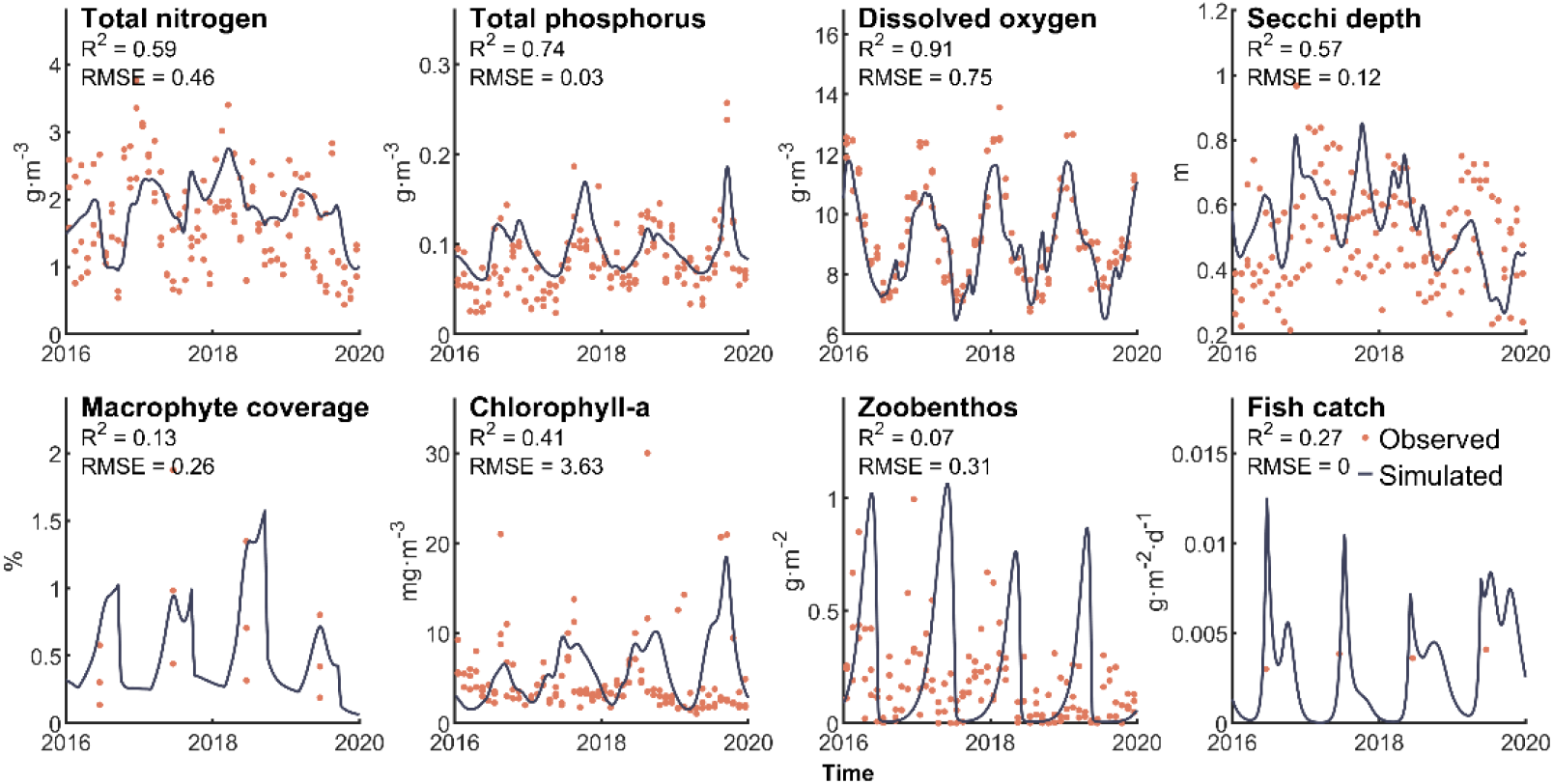
The result of PCLake model calibration between 2016 and 2020. Monthly observational data from Lake Hongze comprised total nitrogen, total phosphorus, dissolved oxygen, Secchi depth, chlorophyll-a, and zoobenthos, while macrophyte coverage and fish harvest were recorded annually. Data from different lake zones are presented; however, the coefficient of determination (R^2^) and root mean square error (RMSE) were calculated based on whole-lake averages and corresponding model simulations.

### 3.2 Hindcast on Fishing Ban and Trophic Cascade

The calibrated PCLake model was applied to conduct a hindcast analysis for the period 2020– 2024 (Figure 3A). Under the fishing ban scenario (FB), total fish biomass and chlorophyll-a showed an increasing trend, while macrophyte coverage declined slightly; other variables remained largely stable. Compared with the no-fishing-ban scenario (no FB), the FB scenario resulted in higher levels of total nitrogen, total phosphorus, chlorophyll-a, and total fish biomass, but lower values of Secchi depth, macrophyte coverage, zoobenthos, and zooplankton.

**Figure 3.**
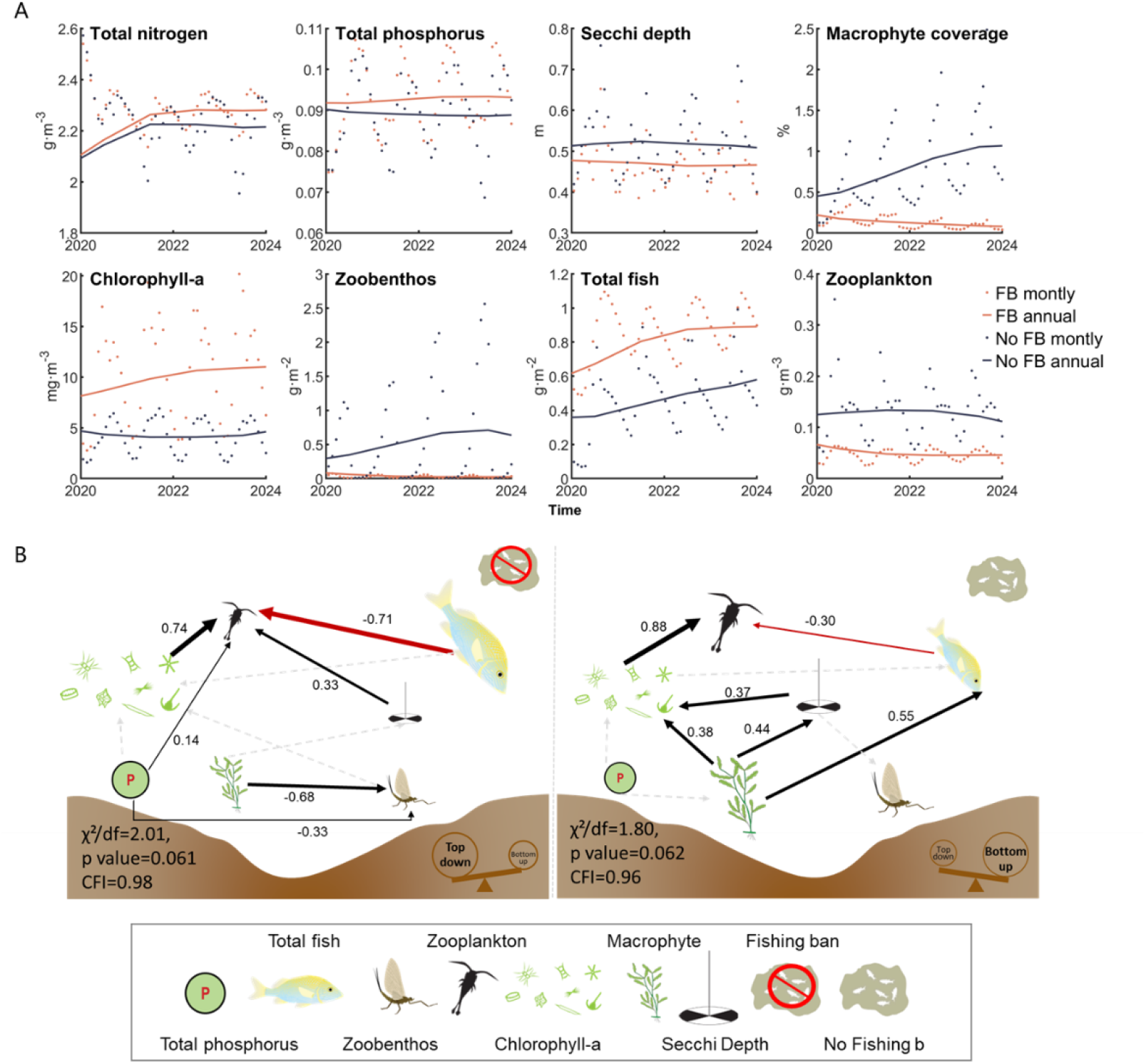
The result of PCLake hindcast and SEM path analysis between 2020 and 2024. (A) “FB” denotes the control scenario with a full fishing ban (fishing intensity = 0), while “no FB” represents the scenario without fishing restrictions. Results are presented both monthly and annually. (B) Two SEM analyses were performed using monthly data from seven variables. Symbols representing these variables were sourced from https://ian.umces.edu/media-library/. Both SEM models exhibited acceptable fit according to the chi-square/degrees of freedom ratio (χ^2^/df), and revealed contrasting dominant trophic cascades under fishing ban (left) and no ban (right) conditions. Red lines indicate negative relationships, black lines positive relationships, with line width corresponding to effect strength. Non-significant paths are shown as dashed grey lines.

Monthly hindcast outputs were further analyzed using structural equation modeling (SEM) to quantify trophic interactions (Figure 3B). Both SEM models exhibited acceptable fit (χ^2^/df < 3, *p* >0.05, CFI>0.9). Monthly hindcast outputs were further analyzed using structural equation modeling (SEM) to quantify trophic interactions (Figure 3B). Both SEM models exhibited acceptable fit (χ^2^/df < 3, p >0.05, CFI>0.9). Without the fishing ban, the path coefficients ranged from –0.71 to 0.74, with strong bottom-up effects as indicated by Chla → Zoo (0.880, p < 0.001) and Veg → Chla (0.378, p < 0.01), while top-down effects were relatively weak (Fish → Zoo: – 0.299, p < 0.001). In contrast, under the fishing ban, top-down control intensified substantially (Fish → Zoo: –0.710, p < 0.001), while bottom-up effects remained strong (Chla → Zoo: 0.740, p < 0.001). In summary, the fishing ban strengthened top-down regulation by fish on zooplankton, while bottom-up control by algae on zooplankton persisted across both scenarios, indicating that the ban shifted the system toward stronger consumer control without eliminating resource-driven effects.

### 3.3 Forecast Under Potential Managements

To evaluate potential regulatory effects from 2025 to 2030, a scenario analysis was conducted using the Hongze Lake PCLake model. Relative changes in annual outputs by 2030 revealed distinct response patterns across scenarios compared to the control (NP- and F0). Variables in group A showed greater sensitivity to nutrient control (differences between NP- and NP--) than to fishing intensity. In contrast, variables in group B showed a clear horizontal gradient across fishing scenarios (F0, F0/1, F0.5, F1), while little difference was observed between the two nutrient reduction levels, suggesting that fishing pressure was the dominant driver. Chlorophyll-a displayed discernible variation along both nutrient and fishing gradients.

Stricter nutrient control significantly reduced nutrient concentrations and increased oxygen levels, whereas fishing bans had minimal impact on these variables. Chlorophyll-a also decreased under lower external loading. Compared to continued fishing ban (F0), scenario F1 (no ban) resulted in markedly lower total fish biomass and chlorophyll-a, but higher zooplankton, zoobenthos, and macrophyte levels. Intermediate fishing scenarios (F0/1 and F0.5) led to moderate effects, with F0.5 yielding slightly lower fish stock and chlorophyll-a, yet higher macrophyte coverage than F0/1. Moreover, differences between F0 and F1 were amplified under stricter nutrient control for chlorophyll-a and all group B variables except for zooplankton.

## 4. Discussion

### 4.1 The Past and Potential antagonistic Effects of Nutrient Control and Fishing Ban

Despite the well-acknowledged negative effect of zooplankivorous fish stocking on water quality, such impact remains neglected in the combination of fishing ban and nutrient control. This neglect was mainly due to the difficulties in conducting process evaluations and the separation of the effectiveness evaluation system. Here, we sought to fill the gap by combining a process-based modeling and path analysis approach in a lake with consistent observations. We confirmed the hypotheses that fishing ban can trigger enhanced top-down control to compromise the effectiveness of nutrient loading control on lake water quality and such antagonism would continue until 2030 if fishing ban will be implemented as planned. Thus, we anticipate that, in the context of comprehensive management, eutrophication restoration would slow down due to more enhanced trophic cascade from fish.

One advantage of combining the PCLake model and SEM analysis is to disentangle both the independent contributions and the indirect empirically supported pathways pathways of combined managements in lake foodweb.

The model revealed divergent trends in lake variables under fishing ban (FB) and non-ban (no FB) conditions (Figure 3). From 2020 to 2024, the FB scenario was characterized by increases in fish biomass and chlorophyll-a concentration. These findings align with reported rises in fish biodiversity and phytoplankton density by Zhang et al. (2024). The expansion of fish stocks resulted from the cessation of fishing, particularly under sufficient availability of zooplankton and zoobenthos as food sources. This also implies that the likelihood of fishing bans leading to an increase in predatory fish populations and a decrease in other fish populations is low. Although macrophytes could offer refuge for zooplankton against predation, their limited coverage— remaining consistently low—rendered this effect negligible. Consequently, zooplankton declined slightly, reducing grazing pressure and contributing to phytoplankton increase. Even under no FB, a modest rise in fish biomass was simulated, attributable to the preexisting seasonal fishing ban (February–June annually since 2012) (Liu 2015), which had already lowered baseline fishing pressure. Continued implementation of such partial bans could enable gradual fish recovery, as reflected in the hindcast. Zoobenthos also increased, likely due to greater stocking of zooplanktivorous over benthivorous fish. Notably, studies such as Ren et al. (2022) indicate that benthic fish can promote sediment nutrient release, stimulating phytoplankton and periphyton growth, which in turn shades and suppresses submerged macrophytes. In our hindcast, Secchi depth remained largely stable despite correlations with other variables, suggesting that factors such as hydrodynamics and mining activities—exacerbated by the lake’s shallow depth and short retention time (Li et al., 2019)—may exert stronger influence on light penetration and macrophyte development.

Our simulations also demonstrate that fishing bans can intensify top-down control, thereby undermining the effectiveness of nutrient reduction measures in improving water quality—an effect projected to persist until 2030 if the ban continues as scheduled (Figure 4). Phytoplankton, positioned at the interface of bottom-up nutrient effects and top-down predation pressure (Figure 3), exhibits complex responses under combined management regimes (Figure 4). Our results indicate that stringent nutrient control (NP--) combined with maintained fishing activity (F1) could optimize water quality by achieving the lowest nutrient and phytoplankton levels alongside the highest transparency, macrophyte coverage, and zoobenthos biomass (Figure 4). However, this scenario also resulted in the lowest fish stock, illustrating a classic ecosystem service trade-off (Rodríguez et al., 2006, Xu et al., 2017). Partial fishing restrictions (e.g., biennial F0/1 or reduced effort F0.5) also attenuated the benefits of nutrient control, reflecting intermediate ecological outcomes. This may result from rapid fish stock expansion during initial ban years, leading to structural surplus (Zhang et al., 2024)—characterized by an overabundance of small-sized or low-value species and an imbalanced size or trophic structure, which may undermine ecosystem stability and complicate post-ban recovery outcomes.

**Figure 4.**
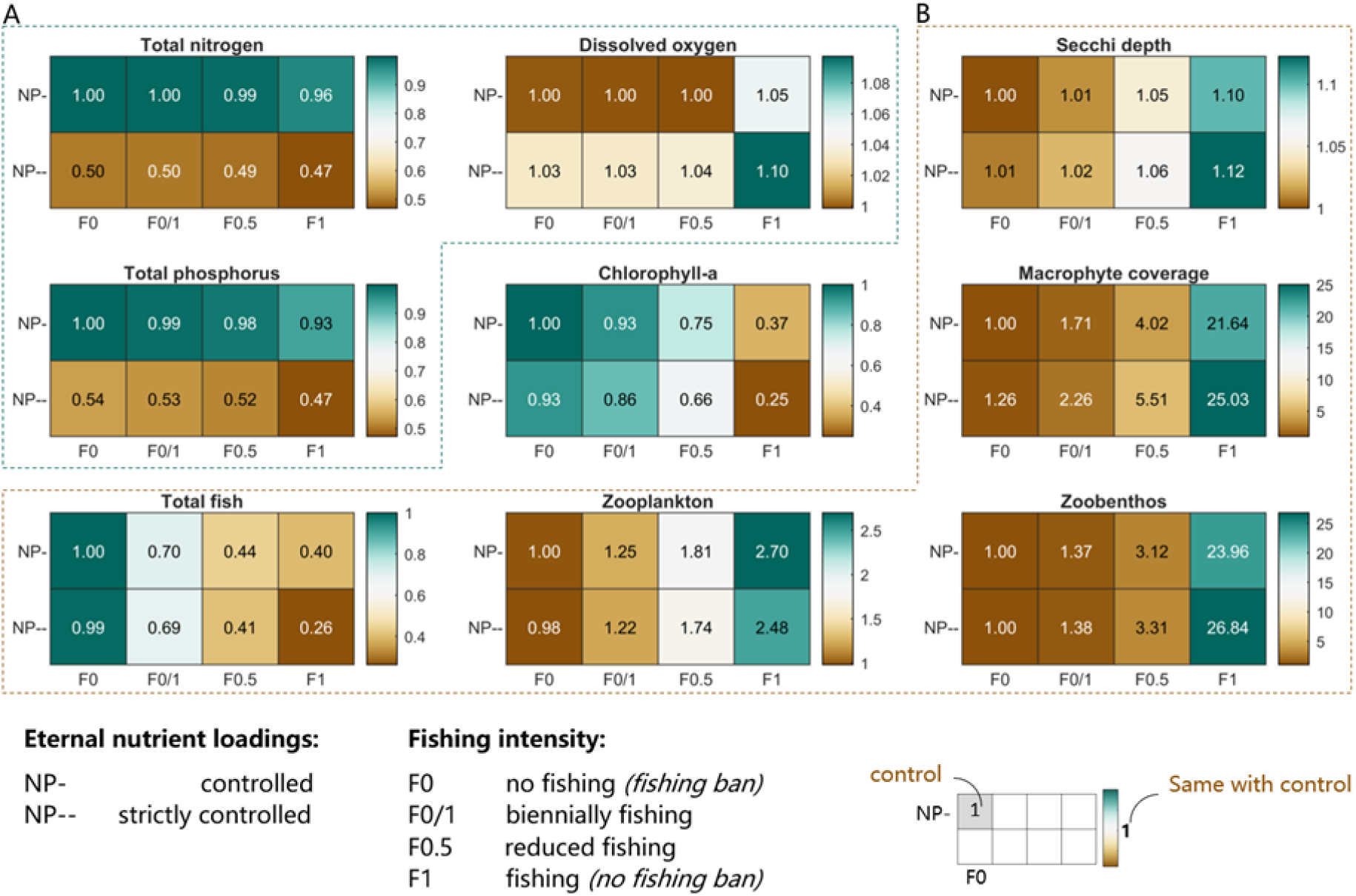
The result of PCLake forecast between 2025 and 2030. The eight scenarios resulted from combining two external nutrient loading conditions (controlled, NP-; strictly controlled, NP--) with four fishing regimes (continuous ban, F0; biennial ban, F0/1; reduced effort, F0.5; no ban, F1). The color bar indicates the relative change compared to the control scenario (NP- and F0), with values above 1 denoting an increase relative to control. Variables are grouped into (A) and (B) based on similar response patterns across scenarios.

Søndergaard et al. (2005) documented an early recovery phase in 12 Danish lakes, with mean inlet phosphorus concentrations declining from 0.56 to 0.13 mg·L-1 in shallow lakes and from 0.27 to 0.12 mg·L-1 in deep lakes, corresponding to a 40%–75% nutrient reduction that may be sufficient to trigger a regime shift from a turbid, algal-dominated to a clear, macrophyte-dominated state. Although Lake Hongze is nearing this critical threshold (Zhang et al., 2024) and both external and internal nutrient levels have declined in recent years, no significant macrophyte recovery has been observed yet. The lake is not yet in a full regime shift, but the intensified trophic cascade from fish may slow down the recovery process once the threshold is crossed. With adjustments to the duration and intensity of fishing bans, the antagonistic effects of enhanced top-down control (from fish) on bottom-up nutrient reduction could be amplified under more stringent nutrient control scenarios (Figure 4). Therefore, complementary strategies—such as reconnecting migratory pathways, restoring habitats, and targeted species stocking (Wang et al., 2022b, Mao et al., 2020, Feng et al., 2023)—should be integrated to simultaneously support biodiversity and eutrophication mitigation.

### 4.2 Evaluations and Limitations of the Modeling

Based on model performance during calibration (2016–2020) and hindcasting (2020–2024), we conclude that the boundary conditions and parameter values of the PCLake model were appropriately configured. The calibrated simulation captured both the magnitude (RMSE) and seasonal dynamics (R^2^) of observed variables comparably with other studies (Figure 2; Table S3). Relatively low R^2^ values for zoobenthos and chlorophyll-a may be attributed to insufficient zooplankton data, leading to uncertain estimates of grazing pressure on phytoplankton and zoobenthos food availability. Although biological variables showed lower fit than water quality parameters, the modified model exhibited improved performance over the original version (Figure S5; Table S3). Measured data from 2020–2023 show that Lake Hongze had TN of 1.4–2.2 mg/L, TP of 0.07–0.10 mg/L (Zhang et al., 2024), Secchi depth of 0.3–0.5 m, and submerged macrophyte coverage near zero (Huang et al., 2025). Wang et al. (2022a) estimated that the fishing ban policy would restore fish resources by up to 30%. Our 2020–2024 hindcast simulations under the fishing ban produced comparable results: TN at 2.25 mg/L (slightly above the measured range), TP at 0.092 mg/L (within the measured range), Secchi depth at 0.47 m (within the measured range), macrophyte coverage declining from 0.2 to 0.05 (approaching zero), and fish biomass rising from 0.6 to 0.9 g/m^2^ (approximately 50% increase, exceeding the 30% projection). These results generally match the field observations, with the minor TN overestimation likely due to the lack of assimilated observational external loading data. Structural equation modeling (SEM) was employed for trophic cascade analysis. According to Schermelleh-Engel & Moosbrugger (2003), a χ^2^/df ratio < 3 indicates good model-data fit. The two SEM models in our study (Figure 3B) both achieved values below this benchmark. Potential sources of uncertainty include simplification of model structure, assumption of fixed pathways and directions—which may vary temporally—and possible nonlinear relationships among variables not captured in the SEM.

The limitations of this study primarily concern the absence of uncertainty analysis and the generalizability of the results. First, uncertainties stemming from the PCLake model structure and the selection of forcing functions for the control scenarios were not quantitatively evaluated. Although originally designed for temperate lakes (Janse 2005), the model for Lake Hongze was adapted to represent omnivorous fish indirectly by incorporating zooplanktivorous fish and adjusting their balance with benthivorous species. Nevertheless, the explicit absence of true omnivorous fish may reduce the reliability of simulations, particularly for biological variables. Second, while the choice of forcing functions—such as average nutrient loading from 2020 to 2024—likely preserves relative differences between scenarios, it may affect the absolute values of predicted variables. Third, the mismatch in temporal scales between lower and higher trophic levels in process-based modeling could influence the simulated responses of certain functional groups. PCLake assumes all trophic levels respond at similar rates, but phytoplankton cycle within days whereas fish dynamics unfold over years—a mismatch that may bias recovery projections and mask delayed or nonlinear ecosystem responses. Additionally, although a mutually antagonistic effect between fishing bans and nutrient reduction was identified in Lake Hongze, synergistic outcomes might occur in lakes dominated by piscivorous fish.

### 4.3 Implications

Our findings hold relevance for a broad range of lakes globally. First, shallow freshwater lakes with food web structures comparable to Lake Hongze are widespread. Many of these systems experience severe eutrophication and water quality degradation, primarily due to pollution and resource overexploitation. The quantitative insights provided here regarding the contributions of different management strategies can support more informed decision-making. Although fishing bans are not traditionally classified as biomanipulation measures, they represent a direct policy approach to mitigating biodiversity loss by addressing key anthropogenic pressures (Wang et al., 2022a). Such targeted interventions, aimed at eliminating specific local stressors, are increasingly relevant in the context of achieving Sustainable Development Goals (SDGs). While the specific antagonistic effects identified here may be most applicable to lakes resembling Lake Hongze in physical and biological characteristics, the proposed research framework offers a transferrable methodology for optimizing net benefits from multiple environmental policies. Importantly, the integration of process-based modeling and path analysis enables simultaneous evaluation of management outcomes and underlying mechanisms—an approach especially critical under future climate change.

Our modeling approach reveals the presence and mechanisms of antagonistic effects between external nutrient control and fishing bans in the restoration of eutrophic freshwater lakes. We project that this antagonism will persist until 2030 if the fishing ban continues as planned, and may intensify under stricter nutrient reduction. Within integrated management frameworks, eutrophication mitigation is likely to slow down due to strengthened trophic cascades originating from fish populations. These results highlight the need for complementary strategies that simultaneously support biodiversity recovery and water quality improvement. Furthermore, management combinations should be implemented within systematic frameworks—featuring quantifiable feedback mechanisms—to address inherent trade-offs among ecosystem services. This study also demonstrates how integrating process-based modeling with path analysis can help quantify the contributions and underlying processes of management measures with complex interactions. Future improvements in model precision are necessary to enhance the reliability of strategies aimed at maximizing net benefits from multi-objective environmental policies. Improved model performance and credibility require more comprehensive observation data, emphasizing the importance of sustained monitoring of lake water quality and ecosystem status.

## Acknowledgements

This work was supported by the Joint Funds of the National Natural Science Foundation of China (No. U24A20639, 42171119, 42507663), and the State Key Laboratory of Lake and Watershed Science for Water Security (NKL2023-ZD02).

## Data and code availability

The code of the original model used in this study is available at https://github.com/pcmodel/PCModel. The data and code for model analysis and result visualization of this study are available in the Supporting Information.

## References

APHA (2012) Standard Methods for the Examination of Water and Wastewater. American Public Health Association, Washington, DC, 22nd edition.

Bennett, N. D., B. F. W. Croke, G. Guariso, J. H. A. Guillaume, S. H. Hamilton, A. J. Jakeman, S. Marsili-Libelli, L. T. H. Newham, J. P. Norton, C. Perrin, S. A. Pierce, B. Robson, R. Seppelt, A. A. Voinov, B. D. Fath & V. Andreassian (2013) Characterising performance of environmental models. Environmental Modelling & Software, 40, 1–20. 10.1016/j.envsoft.2012.09.011

Birk, S., D. Chapman, L. Carvalho, B. M. Spears, H. E. Andersen, C. Argillier, S. Auer, A. Baattrup-Pedersen, L. Banin, M. Beklioğlu, E. Bondar-Kunze, A. Borja, P. Branco, T. Bucak, A. D. Buijse, A. C. Cardoso, R.-M. Couture, F. Cremona, D. de Zwart, C. K. Feld, M. T. Ferreira, H. Feuchtmayr, M. O. Gessner, A. Gieswein, L. Globevnik, D. Graeber, W. Graf, C. Gutiérrez-Cánovas, J. Hanganu, U. Işkın, M. Järvinen, E. Jeppesen, N. Kotamäki, M. Kuijper, J. U. Lemm, S. Lu, A. L. Solheim, U. Mischke, S. J. Moe, P. Nõges, T. Nõges, S. J. Ormerod, Y. Panagopoulos, G. Phillips, L. Posthuma, S. Pouso, C. Prudhomme, K. Rankinen, J. J. Rasmussen, J. Richardson, A. Sagouis, J. M. Santos, R. B. Schäfer, R. Schinegger, S. Schmutz, S. C. Schneider, L. Schülting, P. Segurado, K. Stefanidis, B. Sures, S. J. Thackeray, J. Turunen, M. C. Uyarra, M. Venohr, P. C. von der Ohe, N. Willby & D. Hering (2020) Impacts of multiple stressors on freshwater biota across spatial scales and ecosystems. Nature Ecology & Evolution, 4, 1060–1068. 10.1038/s41559-020-1216-4

Cai, Y., Y. Zhang, Z. Wu, Y. Chen, J. Xu & Z. Gong (2016a) Composition, diversity, and environmental correlates of benthic macroinvertebrate communities in the five largest freshwater lakes of China. Hydrobiologia, 788, 85–98. 10.1007/s10750-016-2989-y

Cai, Y. J., Y. J. Lu, J. S. Liu, X. L. Dai, H. Xu, Y. Lu & Z. J. Gong (2016b) Macrozoobenthic community structure in a large shallow lake: Disentangling the effect of eutrophication and wind-wave disturbance. Limnologica, 59, 1–9. 10.1016/j.limno.2016.03.006

Chen, X., J. Liang, L. Zeng, Y. Cao & M. A. Stevenson (2022) Heterogeneity in diatom diversity response to decadal scale eutrophication in floodplain lakes of the middle Yangtze reaches. Journal of Environmental Management, 322. 10.1016/j.jenvman.2022.116164

Cui, J. Y., R. Guo, X. W. Song, Y. Zhang, C. Chen, X. Lv, Y., & Y. Dong, Y., (2021) Spatio-temporal variations of total nitrogen and total phosphorus in lake and inflow/outflow rivers of Lake Hongze, 2010-2019 (in Chinese). Journal of Lake Science, 33(6): 1727–1741.

Feng, K., W. Deng, H. Li, Q. Guo, K. Tao, J. Yuan, J. Liu, Z. Li, S. Lek, B. Hugueny & Q. Wang (2023) Direct and indirect effects of a fishing ban on lacustrine fish community do not result in a full recovery. Journal of Applied Ecology, 60, 2210–2222. 10.1111/1365-2664.14491

Grace, J. (2006) Structural Equation Modeling Natural Systems. Cambridge University Press. 10.1017/CBO9780511617799

Gsell, A.S., Teurlincx, S., Adrian, R., Janssen, A.B.G. (2023) Timing matters: Sampling frequency for early-warning indicators across food web components in a virtual lake. Ecological Indicators 153, 110424. 10.1016/j.ecolind.2023.110424

Hilt, S., M. M. Alirangues Nuñez, E. S. Bakker, I. Blindow, T. A. Davidson, M. Gillefalk, L.-A. Hansson, J. H. Janse, A. B. G. Janssen, E. Jeppesen, T. Kabus, A. Kelly, J. Köhler, T. L. Lauridsen, W. M. Mooij, R. Noordhuis, G. Phillips, J. Rücker, H.-H. Schuster, M. Søndergaard, S. Teurlincx, K. van de Weyer, E. van Donk, A. Waterstraat, N. Willby & C. D. Sayer (2018) Response of Submerged Macrophyte Communities to External and Internal Restoration Measures in North Temperate Shallow Lakes. Frontiers in Plant Science, 9. 10.3389/fpls.2018.00194

Ho, J. C., A. M. Michalak & N. Pahlevan (2019) Widespread global increase in intense lake phytoplankton blooms since the 1980s. Nature, 574, 667–+. 10.1038/s41586-019-1648-7

Hu, M., R. Ma, K. Xue, Z. Cao, J. Xiong, S. A. Loiselle, M. Shen & X. Hou (2024) Eutrophication evolution of lakes in China: Four decades of observations from space. Journal of Hazardous Materials, 470. 10.1016/j.jhazmat.2024.134225

Huang, X., Q. Xu, B. Zhang, C. Kong, L. Fang, X. Gao, L. Sun, L. Li & X. Gong (2025a) An Assessment of the Population Structure and Stock Dynamics of Megalobrama skolkovii During the Early Phase of the Fishing Ban in the Poyang Lake Basin. Fishes, 10. 10.3390/fishes10080378

Huang, Z. H., H. Z. Liu, Y. Q. Wu, R. H. Ma & X. M. Lu (2025b) The spatiotemporal evolution law of key parameters of water environment in typical lakes in the Jianghuai Basin (in Chinese). Jiangsu Water Resources, (05): 1–5.

IBM, C. (2012) IBM SPSS Statistics for Windows, Version 21.0. Armonk, NY: IBM Corp.

Janse, J. H. (2005) Model studies on the eutrophication of shallow lakes and ditches. Wageningen University Wageningen, Ph. D. Dissertation.

Janse, J. H., M. Scheffer, L. Lijklema, L. Van Liere, J. S. Sloot & W. M. Mooij (2010) Estimating the critical phosphorus loading of shallow lakes with the ecosystem model PCLake: Sensitivity, calibration and uncertainty. Ecological Modelling, 221, 654–665. 10.1016/j.ecolmodel.2009.07.023

Janssen, A. B. G., G. B. Arhonditsis, A. Beusen, K. Bolding, L. Bruce, J. Bruggeman, R. M. Couture, A. S. Downing, J. A. Elliott, M. A. Frassl, G. Gal, D. J. Gerla, M. R. Hipsey, F. J. Hu, S. C. Ives, J. H. Janse, E. Jeppesen, K. D. Johnk, D. Kneis, X. Z. Kong, J. J. Kuiper, M. K. Lehmann, C. Lemmen, D. Ozkundakci, T. Petzoldt, K. Rinke, B. J. Robson, R. Sachse, S. A. Schep, M. Schmid, H. Scholten, S. Teurlincx, D. Trolle, T. A. Troost, A. A. Van Dam, L. P. A. Van Gerven, M. Weijerman, S. A. Wells & W. M. Mooij (2015) Exploring, exploiting and evolving diversity of aquatic ecosystem models: a community perspective. Aquatic Ecology, 49, 513–548. 10.1007/s10452-015-9544-1

Janssen, A. B. G., B. Droppers, X. Kong, S. Teurlincx, Y. Tong & C. Kroeze (2021) Characterizing 19 thousand Chinese lakes, ponds and reservoirs by morphometric, climate and sediment characteristics. Water Research, 202. 10.1016/j.watres.2021.117427

Janssen, A. B. G., S. Teurlincx, S. Q. An, J. H. Janse, H. W. Paerl & W. M. Mooij (2014) Alternative stable states in large shallow lakes? Journal of Great Lakes Research, 40, 813–826. 10.1016/j.jglr.2014.09.019

Jenny, J.-P., O. Anneville, F. Arnaud, Y. Baulaz, D. Bouffard, I. Domaizon, S. A. Bocaniov, N. Chèvre, M. Dittrich, J.-M. Dorioz, E. S. Dunlop, G. Dur, J. Guillard, T. Guinaldo, S. Jacquet, A. Jamoneau, Z. Jawed, E. Jeppesen, G. Krantzberg, J. Lenters, B. Leoni, M. Meybeck, V. Nava, T. Nõges, P. Nõges, M. Patelli, V. Pebbles, M.-E. Perga, S. Rasconi, C. R. Ruetz, L. Rudstam, N. Salmaso, S. Sapna, D. Straile, O. Tammeorg, M. R. Twiss, D. G. Uzarski, A.-M. Ventelä, W. F. Vincent, S. W. Wilhelm, S.-Å. Wängberg & G. A. Weyhenmeyer (2020) Scientists’ Warning to Humanity: Rapid degradation of the world’s large lakes. Journal of Great Lakes Research, 46, 686–702. 10.1016/j.jglr.2020.05.006

Jeppesen, E., M. Meerhoff, K. Holmgren, I. Gonzalez-Bergonzoni, F. Teixeira-de Mello, S. A. J. Declerck, L. De Meester, M. Sondergaard, T. L. Lauridsen, R. Bjerring, J. M. Conde-Porcuna, N. Mazzeo, C. Iglesias, M. Reizenstein, H. J. Malmquist, Z. W. Liu, D. Balayla & X. Lazzaro (2010) Impacts of climate warming on lake fish community structure and potential effects on ecosystem function. Hydrobiologia, 646, 73–90. 10.1007/s10750-010-0171-5

Jeppesen, E., M. Sondergaard, M. Meerhoff, T. L. Lauridsen & J. P. Jensen (2007) Shallow lake restoration by nutrient loading reduction - some recent findings and challenges ahead. Hydrobiologia, 584, 239–252. 10.1007/s10750-007-0596-7

Ji, X. M., L. Wen, M. Zhang & F. Yue (2014) Flux analysis of pollutants entering Hongze Lake (in Chinese). Jiangsu Water Resources, (7): 45–46, 48.

Karin Schermelleh-Engel & Helfried Moosbrugger (2003) Evaluating the Fit of Structural Equation Models: Tests of Significance and Descriptive Goodness-of-Fit Measures. Methods of Psychological Research, 8(2): 23–74.

Kuiper, J. J. (2011) Making Eco Logic and Models Work: An Integrative Approach to Lake Ecosystem Modelling. Wageningen University, Wageningen, Ph. D. Dissertation.

Li, B. & P. M. Pu (2003) Study of the Evolution Tendency of Water Quality in Huai River Basin and Hongze Lake (in Chinese). Resources and Environment in the Yangtze Basin, 12(1): 67–73.

Li, N., K. Shi, Y. L. Zhang, Z. J. Gong, K. Peng, Y. B. Zhang & Y. Zha (2019) Decline in Transparency of Lake Hongze from Long-Term MODIS Observations: Possible Causes and Potential Significance. Remote Sensing, 11. ARTN17710.3390/rs11020177

Lin., M. L., T. L. Zhang., S. W. Ye., W. Li., P. Ren., Z. W. Yang., J. S. Liu. & Z. J. Li. (2013) Status of Fish Resources, Historical Variation and Fisheries management strategies in Hongze Lake (in Chinese). ACTA HYDROBIOLOGICA SINICA, 37(6): 1118–1127. 10.7541/2013.152

Liu, X. Z. (2015) Current Situation, Ploblems and Countermeasures of Fishery Resources in Hongze Lake. Nanjing Agricultural University, Nanjing, Ph. D. Dissertation.

Liu, Y. Y., W. Z. Zhang, Y. X. Wang & E. Y. Wang (1979) Economic Fauna of China: Freshwater Mollusca (in Chinese). Science Press, Beijing.

Mao, Z., X. Gu, Y. Cao, M. Zhang, Q. Zeng, H. Chen, R. Shen & E. Jeppesen (2020) The Role of Top-Down and Bottom-Up Control for Phytoplankton in a Subtropical Shallow Eutrophic Lake: Evidence Based on Long-Term Monitoring and Modeling. Ecosystems, 23, 1449–1463. 10.1007/s10021-020-00480-0

Mao, Z. G., X. H. Gu, Z. J. Gong, Q. F. Zeng, H. H. Chen, H. M. Li, S. Y. Zhang & H. Mu (2019) The structure of fish community and changes of fishery resources in Lake Hongze (in Chinese). Journal of Lake Sciences, 31(4): 1109–1119.

Me, W., D. P. Hamilton, C. G. McBride, J. M. Abell & B. J. Hicks (2018) Modelling hydrology and water quality in a mixed land use catchment and eutrophic lake: Effects of nutrient load reductions and climate change. Environmental Modelling & Software, 109, 114–133. 10.1016/j.envsoft.2018.08.001

Meng. F. J., Y. X. Gao, G. D. Xu, Y. M. Zhang, T. Chen, X. W. Mao, J. Chen & Y. J. Jin (2023) Analysis of Nitrogen and Phosphorus Pollution Flux from Huai River into Hongze Lake Based on LOADSET Model and M-K Test. The Administration and Technique of Environmental Monitoring, 35(3): 17–22.

Mooij, W. M., R. J. Brederveld, J. J. M. de Klein, D. L. DeAngelis, A. S. Downing, M. Faber, D. J. Gerla, M. R. Hipsey, J. ‘t Hoen, J. H. Janse, A. B. G. Janssen, M. Jeuken, B. W. Kooi, B. Lischke, T. Petzoldt, L. Postma, S. A. Schep, H. Scholten, S. Teurlincx, C. Thiange, D. Trolle, A. A. van Dam, L. P. A. van Gerven, E. H. van Nes & J. J. Kuiper (2014) Serving many at once: How a database approach can create unity in dynamical ecosystem modelling. Environmental Modelling & Software, 61, 266–273. 10.1016/j.envsoft.2014.04.004

Ni, L., H. Li, L. Zhou, J. Shi, Y. Nie, F. Zhao & S. Li (2023) Structural characteristics of zooplankton communities in Hongze Lake driven by water environmental factors from 2016 to 2020. Environmental Monitoring and Assessment, 195, 1503. 10.1007/s10661-023-12092-x

Qin, B., X. Kong, R. Wang, Y. Zahao & X. Yang (2022) Lake restoration time of Lake Taibai (China): a case study based on paleolimnology and ecosystem modeling. J Paleolimnol, 68, 25–38. 10.1007/s10933-020-00165-7

Qin, B., M. Xu, K. Zhu, Y. Zhao, E. Zhang & R. Wang (2025) Climate warming will significantly affect future restoration and level of ecosystem services in Lake Erhai. Ecological Modelling, 503, 111067. 10.1016/j.ecolmodel.2025.111067

Qin, B. Q., G. Gao, G. W. Zhu, Y. L. Zhang, Y. Z. Song, X. M. Tang, H. Xu & J. M. Deng (2013) Lake eutrophication and its ecosystem response. Chinese Science Bulletin, 58, 961–970. 10.1007/s11434-012-5560-x">

Reed, M. S. (2008) Stakeholder participation for environmental management: A literature review. Biological Conservation, 141, 2417–2431. 10.1016/j.biocon.2008.07.014

Ren, W., Z. Wen, Y. Cao, H. Wang, C. Yuan, X. Zhang, L. Ni, P. Xie, T. Cao, K. Li & E. Jeppesen (2022) Cascading effects of benthic fish impede reinstatement of clear water conditions in lakes: A mesocosm study. Journal of Environmental Management, 301. 10.1016/j.jenvman.2021.113898

Rodríguez, J. P., T. D. Beard, E. M. Bennett, G. S. Cumming, S. J. Cork, J. Agard, A. P. Dobson & G. D. Peterson (2006) Trade-offs across Space, Time, and Ecosystem Services. Ecology and Society, 11.

Rolighed, J., E. Jeppesen, M. Sondergaard, R. Bjerring, J. H. Janse, W. M. Mooij & D. Trolle (2016) Climate Change Will Make Recovery from Eutrophication More Difficult in Shallow Danish Lake Sobygaard. Water, 8. ARTN45910.3390/w8100459

Schindler, D. W. (2012) The dilemma of controlling cultural eutrophication of lakes. Proceedings of the Royal Society B-Biological Sciences, 279, 4322–4333. 10.1098/rspb.2012.1032

Sinha, E., A. M. Michalak & V. Balaji (2017) Eutrophication will increase during the 21st century as a result of precipitation changes. Science, 357, 405–408. 10.1126/science.aan2409

Søndergaard, M., J. P. Jensen & E. Jeppesen (2005) Seasonal response of nutrients to reduced phosphorus loading in 12 Danish lakes. Freshwater Biology, 50, 1605–1615. 10.1111/j.1365-2427.2005.01412.x

Søndergaard, M., E. Jeppesen, T. L. Lauridsen, C. Skov, E. H. Van Nes, R. Roijackers, E. Lammens & R. Portielje (2007) Lake restoration: successes, failures and long-term effects. Journal of Applied Ecology, 44, 1095–1105. 10.1111/j.1365-2664.2007.01363.x

Søndergaard, M., L. Liboriussen, A. R. Pedersen & E. Jeppesen (2008) Lake Restoration by Fish Removal: Short- and Long-Term Effects in 36 Danish Lakes. Ecosystems, 11, 1291–1305. 10.1007/s10021-008-9193-5

Su, H. J., R. Wang, Y. H. Feng, Y. L. Li, Y. Li, J. Chen, C. Xu, S. P. Wang, J. Y. Fang & P. Xie (2020) Long-term empirical evidence, early warning signals and multiple drivers of regime shifts in a lake ecosystem. Journal of Ecology, 109, 3182–3194. 10.1111/1365-2745.13544

Tang, H. Q. (2006) Biosystematic Study on the Chironomid Larvae in China (Diptera: Chironomidae) (Abstract in English). Nankai University, TianJing, Ph. D. Dissertation.

Tigli, M., M. P. Bak, J. H. Janse, M. Strokal & A. B. G. Janssen (2024) The future of algal blooms in lakes globally is in our hands. Water Res, 268, 122533. 10.1016/j.watres.2024.122533

Wang, H., P. Wang, C. Xu, Y. Sun, L. Shi, L. Zhou, E. Jeppesen, J. Chen & P. Xie (2022a) Can the “10-year fishing ban” rescue biodiversity of the Yangtze River? Innovation (Cambridge (Mass.)), 3, 100235. 10.1016/j.xinn.2022.100235">

Wang, R., C. P. Doncaster, W. Zheng, M. Xu, H. Yang, Y. Li, Y. Cai, Y. Zhao, E. Zhang, X. Yang & B. Qin (2023) High phytoplankton diversity in eutrophic states impedes lake recovery. Journal of Biogeography, 50, 1914–1925. 10.1111/jbi.14698

Wang, R., Y. Han, F. Fan, J. Molinos, J. Xu, K. Wang, D. Wang & Z. Mei (2022b) Need to shift in river-lake connection scheme under the “ten-year fishing ban” in the Yangtze River, China. Ecological Indicators, 143, 109434. 10.1016/j.ecolind.2022.109434

Wu, D., Cao, M., Gao, W., Duan, Z., Zhang, Y. (2024) Simulating critical nutrient loadings of regime shift in the shallow plateau Lake Dianchi. Ecol. Model. 491, 110689. 10.1016/j.ecolmodel.2024.110689

Xu, M., X. H. Dong, X. D. Yang, R. Wang, K. Zhang, Y. J. Zhao, T. A. Davidson & E. Jeppesen (2017) Using palaeolimnological data and historical records to assess long-term dynamics of ecosystem services in typical Yangtze shallow lakes (China). Science of the Total Environment, 584, 791–802. 10.1016/j.scitotenv.2017.01.118

Xue, Y., Kong, X., Mao, Z., Zhang, C., Xue, B., Shi, X., Gu, X. (2025) Hydrological variation drives changes in food web structure and ecosystem function with potential hysteresis in a large temperate shallow lake. Journal of Hydrology 650, 132463. 10.1016/j.jhydrol.2024.132463

Yu, C. X., Z. Y. Li, Z. H. Xu & Z. F. Yang (2020) Lake recovery from eutrophication: Quantitative response of trophic states to anthropogenic influences. Ecological Engineering, 143, 105697. ARTN10569710.1016/j.ecoleng.2019.105697

Zhang, X., Yi, Y., Cao, Y., Yang, Z. (2023) Disentangling the effects of phosphorus loading on food web stability in a large shallow lake. J. Environ. Manage. 328, 116991. 10.1016/j.jenvman.2022.116991

Zhang, Y. L., Y. J. Cai, K. Peng, Z. J. Gong, J. H. Luo, Y. Q. Zhou, J. B. Wiei, S. W. He, N. Li & B. Xue (2024) Challenges and protection strategies of ecological environment of lakes along the Eastern Route of the South to North Water Diversion Project (in Chinese). Journal of Lake Sciences, 36 (5), pp.1289–1302.

