## Supplementary material for "The past and potential antagonistic effects of fishing ban against nutrient loading control on eutrophication restoration in Lake Hongze": data and code: sce1.AmosOutput (from another version of this preprint: version 5, 2026-09-09)

sce1.amw


### D:\0 grind\1 HZ\HZpos\Amos\sce1.amw

#### Analysis Summary

#### Date and Time

Date: dinsdag 9 september 2025

Time: 15:00:49

#### Title

sce1: dinsdag 9 september 2025 15:00

#### Groups

#### Group number 1 (Group number 1)

#### Notes for Group (Group number 1)

The model is recursive.

Sample size = 55

#### Variable Summary (Group number 1)

#### Your model contains the following variables (Group number 1)

Observed, endogenous variables

x9

x8

x11

x7

x10

x6

Observed, exogenous variables

x4

Unobserved, exogenous variables

e1

e2

e3

e4

e5

e6

#### Variable counts (Group number 1)

|  |  |
| --- | --- |
| Number of variables in your model: | 13 |
| Number of observed variables: | 7 |
| Number of unobserved variables: | 6 |
| Number of exogenous variables: | 7 |
| Number of endogenous variables: | 6 |

#### Parameter Summary (Group number 1)

|  | Weights | Covariances | Variances | Means | Intercepts | Total |
| --- | --- | --- | --- | --- | --- | --- |
| Fixed | 6 | 0 | 0 | 6 | 0 | 12 |
| Labeled | 0 | 0 | 0 | 0 | 0 | 0 |
| Unlabeled | 10 | 3 | 7 | 1 | 6 | 27 |
| Total | 16 | 3 | 7 | 7 | 6 | 39 |

#### Assessment of normality (Group number 1)

| Variable | min | max | skew | c.r. | kurtosis | c.r. |
| --- | --- | --- | --- | --- | --- | --- |
| x4 | .040 | .173 | 1.629 | 4.931 | 6.898 | 10.443 |
| x8 | 2.771 | 20.189 | .190 | .574 | -1.091 | -1.651 |
| x9 | .006 | .074 | 1.136 | 3.439 | .133 | .201 |
| x6 | .383 | .725 | 1.277 | 3.866 | 1.480 | 2.240 |
| x7 | .043 | .393 | 1.118 | 3.384 | .677 | 1.025 |
| x11 | .026 | .194 | 3.513 | 10.636 | 14.672 | 22.211 |
| x10 | .245 | 1.095 | -.634 | -1.919 | -.250 | -.378 |
| Multivariate |  |  |  |  | 30.765 | 10.163 |

#### Observations farthest from the centroid (Mahalanobis distance) (Group number 1)

| Observation number | Mahalanobis d-squared | p1 | p2 |
| --- | --- | --- | --- |
| 2 | 37.091 | .000 | .000 |
| 1 | 27.419 | .000 | .000 |
| 4 | 23.247 | .002 | .000 |
| 3 | 17.429 | .015 | .009 |
| 5 | 16.400 | .022 | .007 |
| 17 | 13.128 | .069 | .178 |
| 50 | 12.041 | .099 | .303 |
| 10 | 11.757 | .109 | .246 |
| 6 | 11.314 | .125 | .248 |
| 16 | 11.149 | .132 | .184 |
| 11 | 9.500 | .219 | .682 |
| 52 | 8.938 | .257 | .790 |
| 8 | 8.461 | .294 | .861 |
| 26 | 8.402 | .298 | .803 |
| 22 | 7.507 | .378 | .963 |
| 7 | 7.386 | .390 | .952 |
| 51 | 6.829 | .447 | .987 |
| 47 | 6.153 | .522 | .999 |
| 9 | 6.004 | .539 | .999 |
| 20 | 5.890 | .553 | .998 |
| 14 | 5.600 | .587 | .999 |
| 21 | 5.474 | .602 | .999 |
| 29 | 5.400 | .611 | .999 |
| 12 | 5.223 | .633 | .999 |
| 34 | 4.783 | .686 | 1.000 |
| 36 | 4.567 | .713 | 1.000 |
| 38 | 4.546 | .715 | 1.000 |
| 53 | 4.479 | .723 | 1.000 |
| 44 | 4.408 | .732 | 1.000 |
| 46 | 4.335 | .740 | .999 |
| 37 | 4.321 | .742 | .999 |
| 25 | 4.154 | .762 | .999 |
| 30 | 4.095 | .769 | .998 |
| 45 | 4.041 | .775 | .997 |
| 49 | 3.955 | .785 | .996 |
| 35 | 3.868 | .795 | .995 |
| 42 | 3.748 | .808 | .995 |
| 13 | 3.467 | .839 | .998 |
| 55 | 3.400 | .846 | .997 |
| 15 | 3.399 | .846 | .993 |
| 23 | 3.196 | .866 | .995 |
| 32 | 3.186 | .867 | .989 |
| 54 | 3.155 | .870 | .978 |
| 43 | 3.009 | .884 | .978 |
| 31 | 3.003 | .885 | .953 |
| 28 | 2.938 | .891 | .927 |
| 48 | 2.751 | .907 | .934 |
| 19 | 2.745 | .908 | .868 |
| 33 | 2.636 | .916 | .827 |
| 24 | 2.615 | .918 | .707 |
| 41 | 2.597 | .920 | .544 |
| 40 | 2.565 | .922 | .371 |
| 18 | 2.486 | .928 | .234 |
| 27 | 2.435 | .932 | .104 |
| 39 | 2.375 | .936 | .027 |

#### Sample Moments (Group number 1)

#### Sample Covariances (Group number 1)

|  | x4 | x8 | x9 | x6 | x7 | x11 | x10 |
| --- | --- | --- | --- | --- | --- | --- | --- |
| x4 | .000 |  |  |  |  |  |  |
| x8 | .022 | 22.394 |  |  |  |  |  |
| x9 | .000 | -.065 | .000 |  |  |  |  |
| x6 | .000 | -.010 | .000 | .005 |  |  |  |
| x7 | .000 | .137 | -.001 | .003 | .007 |  |  |
| x11 | .000 | .023 | .000 | .001 | .001 | .001 |  |
| x10 | .000 | .490 | -.001 | -.001 | -.001 | -.002 | .041 |

Condition number = 1015801.856

Eigenvalues

22.406 .031 .009 .003 .000 .000 .000

Determinant of sample covariance matrix = .000

#### Sample Correlations (Group number 1)

|  | x4 | x8 | x9 | x6 | x7 | x11 | x10 |
| --- | --- | --- | --- | --- | --- | --- | --- |
| x4 | 1.000 |  |  |  |  |  |  |
| x8 | .227 | 1.000 |  |  |  |  |  |
| x9 | -.299 | -.768 | 1.000 |  |  |  |  |
| x6 | .274 | -.028 | -.352 | 1.000 |  |  |  |
| x7 | -.097 | .348 | -.520 | .525 | 1.000 |  |  |
| x11 | .687 | .182 | -.473 | .486 | .309 | 1.000 |  |
| x10 | .074 | .511 | -.330 | -.061 | -.035 | -.431 | 1.000 |

Condition number = 85.560

Eigenvalues

2.843 1.813 1.229 .716 .244 .122 .033

#### Sample Means (Group number 1)

|  | x4 | x8 | x9 | x6 | x7 | x11 | x10 |
| --- | --- | --- | --- | --- | --- | --- | --- |
|  | .093 | 10.102 | .023 | .478 | .141 | .052 | .807 |

#### Models

#### Default model (Default model)

#### Notes for Model (Default model)

#### Computation of degrees of freedom (Default model)

|  |  |
| --- | --- |
| Number of distinct sample moments: | 35 |
| Number of distinct parameters to be estimated: | 27 |
| Degrees of freedom (35 - 27): | 8 |

#### Result (Default model)

Minimum was achieved

Chi-square = 27.286

Degrees of freedom = 8

Probability level = .001

#### Group number 1 (Group number 1 - Default model)

#### Estimates (Group number 1 - Default model)

#### Scalar Estimates (Group number 1 - Default model)

#### Maximum Likelihood Estimates

#### Regression Weights: (Group number 1 - Default model)

|  |  |  | Estimate | S.E. | C.R. | P | Label |
| --- | --- | --- | --- | --- | --- | --- | --- |
| x8 | <--- | x4 | 52.714 | 31.686 | 1.664 | .096 | par\_2 |
| x9 | <--- | x8 | -.003 | .000 | -8.807 | \*\*\* | par\_1 |
| x6 | <--- | x9 | -3.806 | .721 | -5.281 | \*\*\* | par\_7 |
| x6 | <--- | x8 | -.011 | .003 | -4.217 | \*\*\* | par\_10 |
| x7 | <--- | x6 | .718 | .105 | 6.856 | \*\*\* | par\_8 |
| x7 | <--- | x4 | -1.110 | .465 | -2.385 | .017 | par\_9 |
| x11 | <--- | x7 | .100 | .022 | 4.487 | \*\*\* | par\_3 |
| x11 | <--- | x8 | .000 | .000 | -.506 | .613 | par\_6 |
| x10 | <--- | x11 | -5.537 | .718 | -7.714 | \*\*\* | par\_4 |
| x10 | <--- | x9 | -7.388 | .783 | -9.432 | \*\*\* | par\_5 |

#### Standardized Regression Weights: (Group number 1 - Default model)

|  |  |  | Estimate |
| --- | --- | --- | --- |
| x8 | <--- | x4 | .221 |
| x9 | <--- | x8 | -.768 |
| x6 | <--- | x9 | -.911 |
| x6 | <--- | x8 | -.727 |
| x7 | <--- | x6 | .602 |
| x7 | <--- | x4 | -.249 |
| x11 | <--- | x7 | .348 |
| x11 | <--- | x8 | -.042 |
| x10 | <--- | x11 | -.623 |
| x10 | <--- | x9 | -.581 |

#### Means: (Group number 1 - Default model)

|  |  |  | Estimate | S.E. | C.R. | P | Label |
| --- | --- | --- | --- | --- | --- | --- | --- |
| x4 |  |  | .093 | .003 | 34.520 | \*\*\* | par\_18 |

#### Intercepts: (Group number 1 - Default model)

|  |  |  | Estimate | S.E. | C.R. | P | Label |
| --- | --- | --- | --- | --- | --- | --- | --- |
| x8 |  |  | 5.201 | 3.012 | 1.727 | .084 | par\_16 |
| x9 |  |  | .052 | .004 | 14.367 | \*\*\* | par\_17 |
| x6 |  |  | .682 | .042 | 16.062 | \*\*\* | par\_20 |
| x7 |  |  | -.098 | .067 | -1.472 | .141 | par\_19 |
| x11 |  |  | .041 | .006 | 6.864 | \*\*\* | par\_15 |
| x10 |  |  | 1.269 | .049 | 25.952 | \*\*\* | par\_14 |

#### Covariances: (Group number 1 - Default model)

|  |  |  | Estimate | S.E. | C.R. | P | Label |
| --- | --- | --- | --- | --- | --- | --- | --- |
| e1 | <--> | e2 | .168 | .048 | 3.488 | \*\*\* | par\_13 |
| e4 | <--> | x4 | .000 | .000 | 4.445 | \*\*\* | par\_11 |
| e5 | <--> | x4 | .001 | .000 | 4.546 | \*\*\* | par\_12 |

#### Correlations: (Group number 1 - Default model)

|  |  |  | Estimate |
| --- | --- | --- | --- |
| e1 | <--> | e2 | .539 |
| e4 | <--> | x4 | .741 |
| e5 | <--> | x4 | .538 |

#### Variances: (Group number 1 - Default model)

|  |  |  | Estimate | S.E. | C.R. | P | Label |
| --- | --- | --- | --- | --- | --- | --- | --- |
| x4 |  |  | .000 | .000 | 5.295 | \*\*\* | par\_21 |
| e2 |  |  | 21.238 | 4.087 | 5.196 | \*\*\* | par\_22 |
| e6 |  |  | .000 | .000 | 5.196 | \*\*\* | par\_23 |
| e3 |  |  | .004 | .001 | 5.196 | \*\*\* | par\_24 |
| e1 |  |  | .005 | .001 | 5.195 | \*\*\* | par\_25 |
| e4 |  |  | .001 | .000 | 5.066 | \*\*\* | par\_26 |
| e5 |  |  | .018 | .004 | 5.150 | \*\*\* | par\_27 |

#### Squared Multiple Correlations: (Group number 1 - Default model)

|  |  |  | Estimate |
| --- | --- | --- | --- |
| x8 |  |  | .049 |
| x9 |  |  | .590 |
| x6 |  |  | .341 |
| x7 |  |  | .413 |
| x11 |  |  | -.033 |
| x10 |  |  | .638 |

#### Matrices (Group number 1 - Default model)

#### Implied (for all variables) Covariances (Group number 1 - Default model)

|  | x4 | x8 | x9 | x6 | x7 | x11 | x10 |
| --- | --- | --- | --- | --- | --- | --- | --- |
| x4 | .000 |  |  |  |  |  |  |
| x8 | .021 | 22.326 |  |  |  |  |  |
| x9 | .000 | -.064 | .000 |  |  |  |  |
| x6 | .000 | -.010 | .000 | .005 |  |  |  |
| x7 | .000 | .138 | -.001 | .004 | .008 |  |  |
| x11 | .000 | .029 | .000 | .000 | .000 | .001 |  |
| x10 | .000 | .392 | -.002 | .001 | .002 | -.003 | .051 |

#### Implied (for all variables) Correlations (Group number 1 - Default model)

|  | x4 | x8 | x9 | x6 | x7 | x11 | x10 |
| --- | --- | --- | --- | --- | --- | --- | --- |
| x4 | 1.000 |  |  |  |  |  |  |
| x8 | .221 | 1.000 |  |  |  |  |  |
| x9 | -.170 | -.768 | 1.000 |  |  |  |  |
| x6 | -.006 | -.028 | -.352 | 1.000 |  |  |  |
| x7 | -.252 | .331 | -.480 | .593 | 1.000 |  |  |
| x11 | .657 | .240 | -.262 | .203 | .144 | 1.000 |  |
| x10 | .014 | .368 | -.472 | .076 | .107 | -.502 | 1.000 |

#### Implied (for all variables) Means (Group number 1 - Default model)

|  | x4 | x8 | x9 | x6 | x7 | x11 | x10 |
| --- | --- | --- | --- | --- | --- | --- | --- |
|  | .093 | 10.102 | .023 | .478 | .141 | .052 | .807 |

#### Implied Covariances (Group number 1 - Default model)

|  | x4 | x8 | x9 | x6 | x7 | x11 | x10 |
| --- | --- | --- | --- | --- | --- | --- | --- |
| x4 | .000 |  |  |  |  |  |  |
| x8 | .021 | 22.326 |  |  |  |  |  |
| x9 | .000 | -.064 | .000 |  |  |  |  |
| x6 | .000 | -.010 | .000 | .005 |  |  |  |
| x7 | .000 | .138 | -.001 | .004 | .008 |  |  |
| x11 | .000 | .029 | .000 | .000 | .000 | .001 |  |
| x10 | .000 | .392 | -.002 | .001 | .002 | -.003 | .051 |

#### Implied Correlations (Group number 1 - Default model)

|  | x4 | x8 | x9 | x6 | x7 | x11 | x10 |
| --- | --- | --- | --- | --- | --- | --- | --- |
| x4 | 1.000 |  |  |  |  |  |  |
| x8 | .221 | 1.000 |  |  |  |  |  |
| x9 | -.170 | -.768 | 1.000 |  |  |  |  |
| x6 | -.006 | -.028 | -.352 | 1.000 |  |  |  |
| x7 | -.252 | .331 | -.480 | .593 | 1.000 |  |  |
| x11 | .657 | .240 | -.262 | .203 | .144 | 1.000 |  |
| x10 | .014 | .368 | -.472 | .076 | .107 | -.502 | 1.000 |

#### Implied Means (Group number 1 - Default model)

|  | x4 | x8 | x9 | x6 | x7 | x11 | x10 |
| --- | --- | --- | --- | --- | --- | --- | --- |
|  | .093 | 10.102 | .023 | .478 | .141 | .052 | .807 |

#### Residual Covariances (Group number 1 - Default model)

|  | x4 | x8 | x9 | x6 | x7 | x11 | x10 |
| --- | --- | --- | --- | --- | --- | --- | --- |
| x4 | .000 |  |  |  |  |  |  |
| x8 | .001 | .068 |  |  |  |  |  |
| x9 | .000 | .000 | .000 |  |  |  |  |
| x6 | .000 | .000 | .000 | .000 |  |  |  |
| x7 | .000 | -.001 | .000 | -.001 | -.001 |  |  |
| x11 | .000 | -.005 | .000 | .001 | .000 | .000 |  |
| x10 | .000 | .098 | .001 | -.002 | -.003 | .001 | -.010 |

#### Residual Means (Group number 1 - Default model)

|  | x4 | x8 | x9 | x6 | x7 | x11 | x10 |
| --- | --- | --- | --- | --- | --- | --- | --- |
|  | .000 | .000 | .000 | .000 | .000 | .000 | .000 |

#### Standardized Residual Covariances (Group number 1 - Default model)

|  | x4 | x8 | x9 | x6 | x7 | x11 | x10 |
| --- | --- | --- | --- | --- | --- | --- | --- |
| x4 | .324 |  |  |  |  |  |  |
| x8 | .099 | .016 |  |  |  |  |  |
| x9 | -1.003 | -.014 | .009 |  |  |  |  |
| x6 | 2.124 | -.001 | .000 | .000 |  |  |  |
| x7 | 1.130 | -.024 | -.064 | -.626 | -.600 |  |  |
| x11 | .598 | -.326 | -1.718 | 2.268 | 1.203 | .690 |  |
| x10 | .402 | .633 | 1.162 | -.963 | -.997 | .585 | -.995 |

#### Standardized Residual Means (Group number 1 - Default model)

|  | x4 | x8 | x9 | x6 | x7 | x11 | x10 |
| --- | --- | --- | --- | --- | --- | --- | --- |
|  | .000 | .000 | .000 | .000 | .000 | .000 | .000 |

#### Total Effects (Group number 1 - Default model)

|  | x4 | x8 | x9 | x6 | x7 | x11 |
| --- | --- | --- | --- | --- | --- | --- |
| x8 | 52.714 | .000 | .000 | .000 | .000 | .000 |
| x9 | -.152 | -.003 | .000 | .000 | .000 | .000 |
| x6 | -.023 | .000 | -3.806 | .000 | .000 | .000 |
| x7 | -1.127 | .000 | -2.732 | .718 | .000 | .000 |
| x11 | -.124 | .000 | -.273 | .072 | .100 | .000 |
| x10 | 1.811 | .023 | -5.876 | -.397 | -.553 | -5.537 |

#### Standardized Total Effects (Group number 1 - Default model)

|  | x4 | x8 | x9 | x6 | x7 | x11 |
| --- | --- | --- | --- | --- | --- | --- |
| x8 | .221 | .000 | .000 | .000 | .000 | .000 |
| x9 | -.170 | -.768 | .000 | .000 | .000 | .000 |
| x6 | -.006 | -.028 | -.911 | .000 | .000 | .000 |
| x7 | -.252 | -.017 | -.548 | .602 | .000 | .000 |
| x11 | -.097 | -.048 | -.191 | .210 | .348 | .000 |
| x10 | .159 | .476 | -.462 | -.130 | -.217 | -.623 |

#### Direct Effects (Group number 1 - Default model)

|  | x4 | x8 | x9 | x6 | x7 | x11 |
| --- | --- | --- | --- | --- | --- | --- |
| x8 | 52.714 | .000 | .000 | .000 | .000 | .000 |
| x9 | .000 | -.003 | .000 | .000 | .000 | .000 |
| x6 | .000 | -.011 | -3.806 | .000 | .000 | .000 |
| x7 | -1.110 | .000 | .000 | .718 | .000 | .000 |
| x11 | .000 | .000 | .000 | .000 | .100 | .000 |
| x10 | .000 | .000 | -7.388 | .000 | .000 | -5.537 |

#### Standardized Direct Effects (Group number 1 - Default model)

|  | x4 | x8 | x9 | x6 | x7 | x11 |
| --- | --- | --- | --- | --- | --- | --- |
| x8 | .221 | .000 | .000 | .000 | .000 | .000 |
| x9 | .000 | -.768 | .000 | .000 | .000 | .000 |
| x6 | .000 | -.727 | -.911 | .000 | .000 | .000 |
| x7 | -.249 | .000 | .000 | .602 | .000 | .000 |
| x11 | .000 | -.042 | .000 | .000 | .348 | .000 |
| x10 | .000 | .000 | -.581 | .000 | .000 | -.623 |

#### Indirect Effects (Group number 1 - Default model)

|  | x4 | x8 | x9 | x6 | x7 | x11 |
| --- | --- | --- | --- | --- | --- | --- |
| x8 | .000 | .000 | .000 | .000 | .000 | .000 |
| x9 | -.152 | .000 | .000 | .000 | .000 | .000 |
| x6 | -.023 | .011 | .000 | .000 | .000 | .000 |
| x7 | -.017 | .000 | -2.732 | .000 | .000 | .000 |
| x11 | -.124 | .000 | -.273 | .072 | .000 | .000 |
| x10 | 1.811 | .023 | 1.512 | -.397 | -.553 | .000 |

#### Standardized Indirect Effects (Group number 1 - Default model)

|  | x4 | x8 | x9 | x6 | x7 | x11 |
| --- | --- | --- | --- | --- | --- | --- |
| x8 | .000 | .000 | .000 | .000 | .000 | .000 |
| x9 | -.170 | .000 | .000 | .000 | .000 | .000 |
| x6 | -.006 | .699 | .000 | .000 | .000 | .000 |
| x7 | -.004 | -.017 | -.548 | .000 | .000 | .000 |
| x11 | -.097 | -.006 | -.191 | .210 | .000 | .000 |
| x10 | .159 | .476 | .119 | -.130 | -.217 | .000 |

#### Modification Indices (Group number 1 - Default model)

#### Covariances: (Group number 1 - Default model)

|  |  |  | M.I. | Par Change |
| --- | --- | --- | --- | --- |
| e5 | <--> | e1 | 6.039 | -.002 |

#### Variances: (Group number 1 - Default model)

|  |  |  | M.I. | Par Change |
| --- | --- | --- | --- | --- |

#### Regression Weights: (Group number 1 - Default model)

|  |  |  | M.I. | Par Change |
| --- | --- | --- | --- | --- |

#### Means: (Group number 1 - Default model)

|  |  |  | M.I. | Par Change |
| --- | --- | --- | --- | --- |

#### Intercepts: (Group number 1 - Default model)

|  |  |  | M.I. | Par Change |
| --- | --- | --- | --- | --- |

#### Minimization History (Default model)

| Iteration |  | Negative eigenvalues | Condition # | Smallest eigenvalue | Diameter | F | NTries | Ratio |
| --- | --- | --- | --- | --- | --- | --- | --- | --- |
| 0 | e | 4 |  | -.428 | 9999.000 | 404.031 | 0 | 9999.000 |
| 1 | e\* | 2 |  | -.409 | 1.127 | 217.638 | 18 | .755 |
| 2 | e | 2 |  | -.218 | .188 | 169.217 | 7 | .743 |
| 3 | e\* | 0 | 105138.751 |  | .608 | 114.935 | 7 | .735 |
| 4 | e | 0 | 19286.752 |  | .751 | 92.139 | 6 | .000 |
| 5 | e | 0 | 9427.197 |  | 1.398 | 46.974 | 2 | .000 |
| 6 | e | 0 | 4600.584 |  | .579 | 30.461 | 1 | 1.169 |
| 7 | e | 0 | 3645.395 |  | .112 | 27.692 | 1 | 1.179 |
| 8 | e | 0 | 3259.428 |  | .039 | 27.306 | 1 | 1.129 |
| 9 | e | 0 | 3304.908 |  | .010 | 27.286 | 1 | 1.043 |
| 10 | e | 0 | 3342.970 |  | .001 | 27.286 | 1 | 1.003 |
| 11 | e | 0 | 3311.516 |  | .000 | 27.286 | 1 | 1.000 |

#### Pairwise Parameter Comparisons (Default model)

#### Variance-covariance Matrix of Estimates (Default model)

|  | par\_1 | par\_2 | par\_3 | par\_4 | par\_5 | par\_6 | par\_7 | par\_8 | par\_9 | par\_10 | par\_11 | par\_12 | par\_13 | par\_14 | par\_15 | par\_16 | par\_17 | par\_18 | par\_19 | par\_20 | par\_21 | par\_22 | par\_23 | par\_24 | par\_25 | par\_26 | par\_27 |
| --- | --- | --- | --- | --- | --- | --- | --- | --- | --- | --- | --- | --- | --- | --- | --- | --- | --- | --- | --- | --- | --- | --- | --- | --- | --- | --- | --- |
| par\_1 | .000 |  |  |  |  |  |  |  |  |  |  |  |  |  |  |  |  |  |  |  |  |  |  |  |  |  |  |
| par\_2 | .000 | 1004.019 |  |  |  |  |  |  |  |  |  |  |  |  |  |  |  |  |  |  |  |  |  |  |  |  |  |
| par\_3 | .000 | .000 | .000 |  |  |  |  |  |  |  |  |  |  |  |  |  |  |  |  |  |  |  |  |  |  |  |  |
| par\_4 | .000 | .000 | -.003 | .515 |  |  |  |  |  |  |  |  |  |  |  |  |  |  |  |  |  |  |  |  |  |  |  |
| par\_5 | .000 | .000 | .003 | .124 | .613 |  |  |  |  |  |  |  |  |  |  |  |  |  |  |  |  |  |  |  |  |  |  |
| par\_6 | .000 | .000 | .000 | .000 | .000 | .000 |  |  |  |  |  |  |  |  |  |  |  |  |  |  |  |  |  |  |  |  |  |
| par\_7 | .000 | .000 | .000 | .000 | .000 | .000 | .519 |  |  |  |  |  |  |  |  |  |  |  |  |  |  |  |  |  |  |  |  |
| par\_8 | .000 | .000 | .000 | .000 | .000 | .000 | .000 | .011 |  |  |  |  |  |  |  |  |  |  |  |  |  |  |  |  |  |  |  |
| par\_9 | .000 | 7.954 | .000 | .000 | .000 | .000 | .000 | .000 | .217 |  |  |  |  |  |  |  |  |  |  |  |  |  |  |  |  |  |  |
| par\_10 | .000 | .000 | .000 | .000 | .000 | .000 | .001 | .000 | .000 | .000 |  |  |  |  |  |  |  |  |  |  |  |  |  |  |  |  |  |
| par\_11 | .000 | .000 | .000 | .000 | .000 | .000 | .000 | .000 | .000 | .000 | .000 |  |  |  |  |  |  |  |  |  |  |  |  |  |  |  |  |
| par\_12 | .000 | .000 | .000 | .000 | .000 | .000 | .000 | .000 | .000 | .000 | .000 | .000 |  |  |  |  |  |  |  |  |  |  |  |  |  |  |  |
| par\_13 | .000 | .000 | .000 | .000 | .000 | .000 | .000 | .000 | .000 | .000 | .000 | .000 | .002 |  |  |  |  |  |  |  |  |  |  |  |  |  |  |
| par\_14 | .000 | .000 | .000 | -.030 | -.021 | .000 | .000 | .000 | .000 | .000 | .000 | .000 | .000 | .002 |  |  |  |  |  |  |  |  |  |  |  |  |  |
| par\_15 | .000 | .000 | .000 | .000 | -.002 | .000 | .000 | .000 | .000 | .000 | .000 | .000 | .000 | .000 | .000 |  |  |  |  |  |  |  |  |  |  |  |  |
| par\_16 | .000 | -93.349 | .000 | .000 | .000 | .000 | .000 | .000 | -.739 | .000 | .000 | .000 | .000 | .000 | .000 | 9.072 |  |  |  |  |  |  |  |  |  |  |  |
| par\_17 | .000 | .000 | .000 | .000 | .000 | .000 | .000 | .000 | .000 | .000 | .000 | .000 | .000 | .000 | .000 | .000 | .000 |  |  |  |  |  |  |  |  |  |  |
| par\_18 | .000 | .000 | .000 | .000 | .000 | .000 | .000 | .000 | .000 | .000 | .000 | .000 | .000 | .000 | .000 | .000 | .000 | .000 |  |  |  |  |  |  |  |  |  |
| par\_19 | .000 | -.739 | .000 | .000 | .000 | .000 | .000 | -.005 | -.020 | .000 | .000 | .000 | .000 | .000 | .000 | .072 | .000 | .000 | .004 |  |  |  |  |  |  |  |  |
| par\_20 | .000 | .000 | .000 | .000 | .000 | .000 | -.027 | .000 | .000 | .000 | .000 | .000 | .000 | .000 | .000 | .000 | .000 | .000 | .000 | .002 |  |  |  |  |  |  |  |
| par\_21 | .000 | .000 | .000 | .000 | .000 | .000 | .000 | .000 | .000 | .000 | .000 | .000 | .000 | .000 | .000 | .000 | .000 | .000 | .000 | .000 | .000 |  |  |  |  |  |  |
| par\_22 | .000 | .000 | .000 | .000 | .000 | .000 | .000 | .000 | .000 | .000 | .000 | .000 | .132 | .000 | .000 | .000 | .000 | .000 | .000 | .000 | .000 | 16.705 |  |  |  |  |  |
| par\_23 | .000 | .000 | .000 | .000 | .000 | .000 | .000 | .000 | .000 | .000 | .000 | .000 | .000 | .000 | .000 | .000 | .000 | .000 | .000 | .000 | .000 | .000 | .000 |  |  |  |  |
| par\_24 | .000 | .000 | .000 | .000 | .000 | .000 | .000 | .000 | .000 | .000 | .000 | .000 | .000 | .000 | .000 | .000 | .000 | .000 | .000 | .000 | .000 | .000 | .000 | .000 |  |  |  |
| par\_25 | .000 | .000 | .000 | .000 | .000 | .000 | .000 | .000 | .000 | .000 | .000 | .000 | .000 | .000 | .000 | .000 | .000 | .000 | .000 | .000 | .000 | .001 | .000 | .000 | .000 |  |  |
| par\_26 | .000 | .000 | .000 | .000 | .000 | .000 | .000 | .000 | .000 | .000 | .000 | .000 | .000 | .000 | .000 | .000 | .000 | .000 | .000 | .000 | .000 | .000 | .000 | .000 | .000 | .000 |  |
| par\_27 | .000 | .000 | .000 | .000 | .000 | .000 | .000 | .000 | .000 | .000 | .000 | .000 | .000 | .000 | .000 | .000 | .000 | .000 | .000 | .000 | .000 | .000 | .000 | .000 | .000 | .000 | .000 |

#### Correlations of Estimates (Default model)

|  | par\_1 | par\_2 | par\_3 | par\_4 | par\_5 | par\_6 | par\_7 | par\_8 | par\_9 | par\_10 | par\_11 | par\_12 | par\_13 | par\_14 | par\_15 | par\_16 | par\_17 | par\_18 | par\_19 | par\_20 | par\_21 | par\_22 | par\_23 | par\_24 | par\_25 | par\_26 | par\_27 |
| --- | --- | --- | --- | --- | --- | --- | --- | --- | --- | --- | --- | --- | --- | --- | --- | --- | --- | --- | --- | --- | --- | --- | --- | --- | --- | --- | --- |
| par\_1 | 1.000 |  |  |  |  |  |  |  |  |  |  |  |  |  |  |  |  |  |  |  |  |  |  |  |  |  |  |
| par\_2 | .000 | 1.000 |  |  |  |  |  |  |  |  |  |  |  |  |  |  |  |  |  |  |  |  |  |  |  |  |  |
| par\_3 | .000 | .000 | 1.000 |  |  |  |  |  |  |  |  |  |  |  |  |  |  |  |  |  |  |  |  |  |  |  |  |
| par\_4 | .000 | .000 | -.207 | 1.000 |  |  |  |  |  |  |  |  |  |  |  |  |  |  |  |  |  |  |  |  |  |  |  |
| par\_5 | .000 | .000 | .154 | .221 | 1.000 |  |  |  |  |  |  |  |  |  |  |  |  |  |  |  |  |  |  |  |  |  |  |
| par\_6 | .000 | .000 | -.264 | .062 | .456 | 1.000 |  |  |  |  |  |  |  |  |  |  |  |  |  |  |  |  |  |  |  |  |  |
| par\_7 | .000 | .000 | .000 | .000 | .000 | .000 | 1.000 |  |  |  |  |  |  |  |  |  |  |  |  |  |  |  |  |  |  |  |  |
| par\_8 | .000 | .000 | .000 | .000 | .000 | .000 | .000 | 1.000 |  |  |  |  |  |  |  |  |  |  |  |  |  |  |  |  |  |  |  |
| par\_9 | .000 | .539 | .000 | .000 | .000 | .000 | .000 | .005 | 1.000 |  |  |  |  |  |  |  |  |  |  |  |  |  |  |  |  |  |  |
| par\_10 | .000 | .000 | .000 | .000 | .000 | .000 | .768 | .000 | .000 | 1.000 |  |  |  |  |  |  |  |  |  |  |  |  |  |  |  |  |  |
| par\_11 | .000 | .000 | .011 | .378 | .026 | -.092 | .000 | .000 | .000 | .000 | 1.000 |  |  |  |  |  |  |  |  |  |  |  |  |  |  |  |  |
| par\_12 | .000 | .000 | -.081 | .081 | .095 | .111 | .000 | .000 | .000 | .000 | .014 | 1.000 |  |  |  |  |  |  |  |  |  |  |  |  |  |  |  |
| par\_13 | .000 | .000 | .000 | .000 | .000 | .000 | .000 | .020 | .000 | .000 | .000 | .000 | 1.000 |  |  |  |  |  |  |  |  |  |  |  |  |  |  |
| par\_14 | .000 | .000 | .101 | -.851 | -.543 | -.218 | .000 | .000 | .000 | .000 | -.300 | -.098 | .000 | 1.000 |  |  |  |  |  |  |  |  |  |  |  |  |  |
| par\_15 | .000 | .000 | -.331 | .063 | -.429 | -.620 | .000 | .000 | .000 | .000 | .064 | -.041 | .000 | .112 | 1.000 |  |  |  |  |  |  |  |  |  |  |  |  |
| par\_16 | .000 | -.978 | .000 | .000 | .000 | .000 | .000 | .000 | -.528 | .000 | .000 | .000 | .000 | .000 | .000 | 1.000 |  |  |  |  |  |  |  |  |  |  |  |
| par\_17 | -.906 | .000 | .000 | .000 | .000 | .000 | .000 | .000 | .000 | .000 | .000 | .000 | .000 | .000 | .000 | .000 | 1.000 |  |  |  |  |  |  |  |  |  |  |
| par\_18 | .000 | .000 | .000 | .000 | .000 | .000 | .000 | .000 | .000 | .000 | .000 | .000 | .000 | .203 | .440 | .000 | .000 | 1.000 |  |  |  |  |  |  |  |  |  |
| par\_19 | .000 | -.349 | .000 | .000 | .000 | .000 | .000 | -.750 | -.650 | .000 | .000 | .000 | -.015 | .000 | .000 | .357 | .000 | .000 | 1.000 |  |  |  |  |  |  |  |  |
| par\_20 | .000 | .000 | .000 | .000 | .000 | .000 | -.890 | .000 | .000 | -.948 | .000 | .000 | .000 | .000 | .000 | .000 | .000 | .000 | .000 | 1.000 |  |  |  |  |  |  |  |
| par\_21 | .000 | .000 | -.174 | .535 | .076 | .061 | .000 | .000 | .000 | .000 | .816 | .448 | .000 | -.440 | .046 | .000 | .000 | .000 | .000 | .000 | 1.000 |  |  |  |  |  |  |
| par\_22 | .000 | .000 | .000 | .000 | .000 | .000 | .000 | .000 | .000 | .000 | .000 | .000 | .671 | .000 | .000 | .000 | .000 | .000 | .000 | .000 | .000 | 1.000 |  |  |  |  |  |
| par\_23 | .000 | .000 | .000 | .000 | .000 | .000 | .000 | .000 | .000 | .000 | .000 | .000 | .000 | .000 | .000 | .000 | .000 | .000 | .000 | .000 | .000 | .000 | 1.000 |  |  |  |  |
| par\_24 | .000 | .000 | .000 | .000 | .000 | .000 | .000 | .000 | .000 | .000 | .000 | .000 | .000 | .000 | .000 | .000 | .000 | .000 | .000 | .000 | .000 | .000 | .000 | 1.000 |  |  |  |
| par\_25 | .000 | .000 | .000 | .000 | .000 | .000 | .000 | .018 | .000 | .000 | .000 | .000 | .671 | .000 | .000 | .000 | .000 | .000 | -.013 | .000 | .000 | .291 | .000 | .000 | 1.000 |  |  |
| par\_26 | .000 | .000 | .180 | -.038 | -.039 | -.173 | .000 | .000 | .000 | .000 | .848 | -.027 | .000 | .044 | .036 | .000 | .000 | .000 | .000 | .000 | .513 | .000 | .000 | .000 | .000 | 1.000 |  |
| par\_27 | .000 | .000 | .000 | .093 | .112 | .048 | .000 | .000 | .000 | .000 | .030 | .882 | .000 | -.114 | -.037 | .000 | .000 | .000 | .000 | .000 | .338 | .000 | .000 | .000 | .000 | -.007 | 1.000 |

#### Model Fit Summary

#### CMIN

| Model | NPAR | CMIN | DF | P | CMIN/DF |
| --- | --- | --- | --- | --- | --- |
| Default model | 27 | 27.286 | 8 | .001 | 3.411 |
| Saturated model | 35 | .000 | 0 |  |  |
| Independence model | 14 | 291.938 | 21 | .000 | 13.902 |

#### Baseline Comparisons

| Model | NFI Delta1 | RFI rho1 | IFI Delta2 | TLI rho2 | CFI |
| --- | --- | --- | --- | --- | --- |
| Default model | .907 | .755 | .932 | .813 | .929 |
| Saturated model | 1.000 |  | 1.000 |  | 1.000 |
| Independence model | .000 | .000 | .000 | .000 | .000 |

#### Parsimony-Adjusted Measures

| Model | PRATIO | PNFI | PCFI |
| --- | --- | --- | --- |
| Default model | .381 | .345 | .354 |
| Saturated model | .000 | .000 | .000 |
| Independence model | 1.000 | .000 | .000 |

#### NCP

| Model | NCP | LO 90 | HI 90 |
| --- | --- | --- | --- |
| Default model | 19.286 | 7.046 | 39.106 |
| Saturated model | .000 | .000 | .000 |
| Independence model | 270.938 | 219.422 | 329.899 |

#### FMIN

| Model | FMIN | F0 | LO 90 | HI 90 |
| --- | --- | --- | --- | --- |
| Default model | .505 | .357 | .130 | .724 |
| Saturated model | .000 | .000 | .000 | .000 |
| Independence model | 5.406 | 5.017 | 4.063 | 6.109 |

#### RMSEA

| Model | RMSEA | LO 90 | HI 90 | PCLOSE |
| --- | --- | --- | --- | --- |
| Default model | .211 | .128 | .301 | .002 |
| Independence model | .489 | .440 | .539 | .000 |

#### AIC

| Model | AIC | BCC | BIC | CAIC |
| --- | --- | --- | --- | --- |
| Default model | 81.286 | 90.677 |  |  |
| Saturated model | 70.000 | 82.174 |  |  |
| Independence model | 319.938 | 324.807 |  |  |

#### ECVI

| Model | ECVI | LO 90 | HI 90 | MECVI |
| --- | --- | --- | --- | --- |
| Default model | 1.505 | 1.279 | 1.872 | 1.679 |
| Saturated model | 1.296 | 1.296 | 1.296 | 1.522 |
| Independence model | 5.925 | 4.971 | 7.017 | 6.015 |

#### HOELTER

| Model | HOELTER .05 | HOELTER .01 |
| --- | --- | --- |
| Default model | 31 | 40 |
| Independence model | 7 | 8 |

#### Execution time summary

|  |  |
| --- | --- |
| Minimization: | .022 |
| Miscellaneous: | .202 |
| Bootstrap: | .000 |
| Total: | .224 |
