## Supplementary material for "The past and potential antagonistic effects of fishing ban against nutrient loading control on eutrophication restoration in Lake Hongze": data and code: sce2.AmosOutput (from another version of this preprint: version 5, 2026-09-09)

sce2.amw


### D:\0 grind\1 HZ\HZpos\Amos\sce2.amw

#### Analysis Summary

#### Date and Time

Date: dinsdag 9 september 2025

Time: 15:24:05

#### Title

sce2: dinsdag 9 september 2025 15:24

#### Groups

#### Group number 1 (Group number 1)

#### Notes for Group (Group number 1)

The model is recursive.

Sample size = 55

#### Variable Summary (Group number 1)

#### Your model contains the following variables (Group number 1)

Observed, endogenous variables

x9

x8

x11

x7

x6

Observed, exogenous variables

x4

x10

Unobserved, exogenous variables

e1

e2

e3

e4

e6

#### Variable counts (Group number 1)

|  |  |
| --- | --- |
| Number of variables in your model: | 12 |
| Number of observed variables: | 7 |
| Number of unobserved variables: | 5 |
| Number of exogenous variables: | 7 |
| Number of endogenous variables: | 5 |

#### Parameter Summary (Group number 1)

|  | Weights | Covariances | Variances | Means | Intercepts | Total |
| --- | --- | --- | --- | --- | --- | --- |
| Fixed | 5 | 0 | 0 | 5 | 0 | 10 |
| Labeled | 0 | 0 | 0 | 0 | 0 | 0 |
| Unlabeled | 10 | 3 | 7 | 2 | 5 | 27 |
| Total | 15 | 3 | 7 | 7 | 5 | 37 |

#### Assessment of normality (Group number 1)

| Variable | min | max | skew | c.r. | kurtosis | c.r. |
| --- | --- | --- | --- | --- | --- | --- |
| x4 | .040 | .173 | 1.988 | 6.020 | 8.076 | 12.225 |
| x10 | .069 | .992 | .376 | 1.138 | -.396 | -.600 |
| x8 | 1.589 | 7.373 | -.088 | -.267 | -1.271 | -1.923 |
| x9 | .006 | 2.561 | 1.349 | 4.083 | .689 | 1.042 |
| x6 | .414 | .761 | .826 | 2.502 | .097 | .146 |
| x7 | .111 | 2.483 | 1.124 | 3.404 | .847 | 1.282 |
| x11 | .053 | .350 | 1.035 | 3.133 | 2.197 | 3.326 |
| Multivariate |  |  |  |  | 26.579 | 8.780 |

#### Observations farthest from the centroid (Mahalanobis distance) (Group number 1)

| Observation number | Mahalanobis d-squared | p1 | p2 |
| --- | --- | --- | --- |
| 13 | 31.510 | .000 | .003 |
| 2 | 23.093 | .002 | .004 |
| 1 | 22.777 | .002 | .000 |
| 53 | 22.174 | .002 | .000 |
| 4 | 18.166 | .011 | .000 |
| 52 | 13.375 | .063 | .135 |
| 3 | 13.190 | .068 | .076 |
| 5 | 12.471 | .086 | .098 |
| 41 | 11.022 | .138 | .343 |
| 51 | 10.769 | .149 | .299 |
| 50 | 10.649 | .155 | .223 |
| 16 | 10.095 | .183 | .301 |
| 14 | 8.818 | .266 | .737 |
| 6 | 8.074 | .326 | .901 |
| 15 | 7.966 | .336 | .872 |
| 38 | 7.903 | .341 | .823 |
| 49 | 7.679 | .362 | .830 |
| 29 | 7.059 | .423 | .944 |
| 30 | 6.748 | .456 | .963 |
| 37 | 6.321 | .503 | .987 |
| 26 | 6.272 | .508 | .978 |
| 27 | 6.025 | .537 | .985 |
| 25 | 5.662 | .580 | .995 |
| 48 | 5.397 | .612 | .997 |
| 12 | 5.073 | .651 | .999 |
| 40 | 5.019 | .658 | .998 |
| 39 | 4.854 | .678 | .999 |
| 7 | 4.727 | .693 | .999 |
| 10 | 4.710 | .695 | .997 |
| 36 | 4.657 | .702 | .995 |
| 9 | 4.646 | .703 | .990 |
| 8 | 4.349 | .739 | .996 |
| 47 | 4.226 | .753 | .996 |
| 11 | 4.097 | .769 | .996 |
| 54 | 3.746 | .809 | .999 |
| 46 | 3.617 | .823 | .999 |
| 35 | 3.441 | .841 | .999 |
| 34 | 3.431 | .842 | .999 |
| 17 | 3.382 | .848 | .997 |
| 45 | 3.300 | .856 | .996 |
| 33 | 3.292 | .857 | .991 |
| 28 | 3.224 | .864 | .986 |
| 18 | 3.026 | .883 | .990 |
| 44 | 2.926 | .892 | .987 |
| 32 | 2.708 | .911 | .991 |
| 22 | 2.583 | .921 | .989 |
| 21 | 2.540 | .924 | .978 |
| 31 | 2.528 | .925 | .948 |
| 23 | 2.190 | .949 | .978 |
| 20 | 2.182 | .949 | .940 |
| 24 | 1.953 | .962 | .944 |
| 19 | 1.516 | .982 | .982 |
| 55 | 1.421 | .985 | .949 |
| 43 | 1.365 | .987 | .832 |
| 42 | 1.061 | .994 | .710 |

#### Sample Moments (Group number 1)

#### Sample Covariances (Group number 1)

|  | x4 | x10 | x8 | x9 | x6 | x7 | x11 |
| --- | --- | --- | --- | --- | --- | --- | --- |
| x4 | .000 |  |  |  |  |  |  |
| x10 | .001 | .048 |  |  |  |  |  |
| x8 | .014 | .207 | 2.735 |  |  |  |  |
| x9 | -.004 | -.003 | -.188 | .490 |  |  |  |
| x6 | .000 | .004 | .081 | -.005 | .008 |  |  |
| x7 | .000 | .084 | .494 | .078 | .020 | .278 |  |
| x11 | .000 | .001 | .050 | .015 | .002 | .007 | .003 |

Condition number = 9731.307

Eigenvalues

2.864 .518 .156 .020 .004 .001 .000

Determinant of sample covariance matrix = .000

#### Sample Correlations (Group number 1)

|  | x4 | x10 | x8 | x9 | x6 | x7 | x11 |
| --- | --- | --- | --- | --- | --- | --- | --- |
| x4 | 1.000 |  |  |  |  |  |  |
| x10 | .154 | 1.000 |  |  |  |  |  |
| x8 | .415 | .568 | 1.000 |  |  |  |  |
| x9 | -.288 | -.018 | -.162 | 1.000 |  |  |  |
| x6 | .154 | .225 | .560 | -.084 | 1.000 |  |  |
| x7 | -.030 | .728 | .567 | .213 | .441 | 1.000 |  |
| x11 | .165 | .093 | .544 | .387 | .409 | .239 | 1.000 |

Condition number = 24.201

Eigenvalues

2.880 1.474 1.176 .762 .402 .187 .119

#### Sample Means (Group number 1)

|  | x4 | x10 | x8 | x9 | x6 | x7 | x11 |
| --- | --- | --- | --- | --- | --- | --- | --- |
|  | .089 | .430 | 4.210 | .532 | .532 | .734 | .140 |

#### Models

#### Default model (Default model)

#### Notes for Model (Default model)

#### Computation of degrees of freedom (Default model)

|  |  |
| --- | --- |
| Number of distinct sample moments: | 35 |
| Number of distinct parameters to be estimated: | 27 |
| Degrees of freedom (35 - 27): | 8 |

#### Result (Default model)

Minimum was achieved

Chi-square = 28.851

Degrees of freedom = 8

Probability level = .000

#### Group number 1 (Group number 1 - Default model)

#### Estimates (Group number 1 - Default model)

#### Scalar Estimates (Group number 1 - Default model)

#### Maximum Likelihood Estimates

#### Regression Weights: (Group number 1 - Default model)

|  |  |  | Estimate | S.E. | C.R. | P | Label |
| --- | --- | --- | --- | --- | --- | --- | --- |
| x8 | <--- | x4 | 33.520 | 9.993 | 3.354 | \*\*\* | par\_2 |
| x9 | <--- | x8 | -.121 | .054 | -2.252 | .024 | par\_1 |
| x9 | <--- | x10 | .461 | .428 | 1.076 | .282 | par\_8 |
| x6 | <--- | x8 | .030 | .006 | 4.781 | \*\*\* | par\_7 |
| x6 | <--- | x9 | .001 | .014 | .060 | .952 | par\_13 |
| x7 | <--- | x6 | 1.841 | .452 | 4.069 | \*\*\* | par\_5 |
| x7 | <--- | x4 | -3.235 | 1.889 | -1.713 | .087 | par\_6 |
| x11 | <--- | x7 | .011 | .016 | .679 | .497 | par\_3 |
| x11 | <--- | x8 | .024 | .004 | 6.415 | \*\*\* | par\_4 |
| x11 | <--- | x10 | -.097 | .038 | -2.540 | .011 | par\_9 |

#### Standardized Regression Weights: (Group number 1 - Default model)

|  |  |  | Estimate |
| --- | --- | --- | --- |
| x8 | <--- | x4 | .415 |
| x9 | <--- | x8 | -.275 |
| x9 | <--- | x10 | .139 |
| x6 | <--- | x8 | .561 |
| x6 | <--- | x9 | .007 |
| x7 | <--- | x6 | .321 |
| x7 | <--- | x4 | -.132 |
| x11 | <--- | x7 | .090 |
| x11 | <--- | x8 | .630 |
| x11 | <--- | x10 | -.343 |

#### Means: (Group number 1 - Default model)

|  |  |  | Estimate | S.E. | C.R. | P | Label |
| --- | --- | --- | --- | --- | --- | --- | --- |
| x4 |  |  | .089 | .003 | 31.989 | \*\*\* | par\_18 |
| x10 |  |  | .430 | .030 | 14.367 | \*\*\* | par\_14 |

#### Intercepts: (Group number 1 - Default model)

|  |  |  | Estimate | S.E. | C.R. | P | Label |
| --- | --- | --- | --- | --- | --- | --- | --- |
| x8 |  |  | 1.220 | .915 | 1.334 | .182 | par\_16 |
| x9 |  |  | .843 | .307 | 2.748 | .006 | par\_17 |
| x6 |  |  | .406 | .031 | 13.126 | \*\*\* | par\_20 |
| x7 |  |  | .044 | .273 | .161 | .872 | par\_19 |
| x11 |  |  | .073 | .020 | 3.611 | \*\*\* | par\_15 |

#### Covariances: (Group number 1 - Default model)

|  |  |  | Estimate | S.E. | C.R. | P | Label |
| --- | --- | --- | --- | --- | --- | --- | --- |
| e1 | <--> | x10 | .079 | .018 | 4.404 | \*\*\* | par\_10 |
| e1 | <--> | e6 | .089 | .031 | 2.879 | .004 | par\_12 |
| e4 | <--> | e6 | .019 | .005 | 4.086 | \*\*\* | par\_11 |

#### Correlations: (Group number 1 - Default model)

|  |  |  | Estimate |
| --- | --- | --- | --- |
| e1 | <--> | x10 | .749 |
| e1 | <--> | e6 | .268 |
| e4 | <--> | e6 | .612 |

#### Variances: (Group number 1 - Default model)

|  |  |  | Estimate | S.E. | C.R. | P | Label |
| --- | --- | --- | --- | --- | --- | --- | --- |
| x4 |  |  | .000 | .000 | 5.196 | \*\*\* | par\_21 |
| e2 |  |  | 2.264 | .436 | 5.196 | \*\*\* | par\_22 |
| x10 |  |  | .048 | .009 | 5.196 | \*\*\* | par\_23 |
| e6 |  |  | .480 | .091 | 5.281 | \*\*\* | par\_24 |
| e3 |  |  | .005 | .001 | 5.196 | \*\*\* | par\_25 |
| e1 |  |  | .228 | .044 | 5.196 | \*\*\* | par\_26 |
| e4 |  |  | .002 | .000 | 5.196 | \*\*\* | par\_27 |

#### Squared Multiple Correlations: (Group number 1 - Default model)

|  |  |  | Estimate |
| --- | --- | --- | --- |
| x8 |  |  | .172 |
| x9 |  |  | .095 |
| x6 |  |  | .313 |
| x7 |  |  | .103 |
| x11 |  |  | .494 |

#### Matrices (Group number 1 - Default model)

#### Implied (for all variables) Covariances (Group number 1 - Default model)

|  | x4 | x10 | x8 | x9 | x6 | x7 | x11 |
| --- | --- | --- | --- | --- | --- | --- | --- |
| x4 | .000 |  |  |  |  |  |  |
| x10 | .000 | .048 |  |  |  |  |  |
| x8 | .014 | .000 | 2.735 |  |  |  |  |
| x9 | -.002 | .022 | -.331 | .531 |  |  |  |
| x6 | .000 | .000 | .081 | -.009 | .008 |  |  |
| x7 | -.001 | .079 | .104 | .113 | .013 | .254 |  |
| x11 | .000 | -.004 | .066 | .010 | .002 | -.002 | .004 |

#### Implied (for all variables) Correlations (Group number 1 - Default model)

|  | x4 | x10 | x8 | x9 | x6 | x7 | x11 |
| --- | --- | --- | --- | --- | --- | --- | --- |
| x4 | 1.000 |  |  |  |  |  |  |
| x10 | .000 | 1.000 |  |  |  |  |  |
| x8 | .415 | .000 | 1.000 |  |  |  |  |
| x9 | -.114 | .139 | -.275 | 1.000 |  |  |  |
| x6 | .232 | .001 | .559 | -.147 | 1.000 |  |  |
| x7 | -.057 | .710 | .125 | .308 | .293 | 1.000 |  |
| x11 | .257 | -.279 | .642 | .221 | .381 | -.074 | 1.000 |

#### Implied (for all variables) Means (Group number 1 - Default model)

|  | x4 | x10 | x8 | x9 | x6 | x7 | x11 |
| --- | --- | --- | --- | --- | --- | --- | --- |
|  | .089 | .430 | 4.210 | .532 | .532 | .734 | .140 |

#### Implied Covariances (Group number 1 - Default model)

|  | x4 | x10 | x8 | x9 | x6 | x7 | x11 |
| --- | --- | --- | --- | --- | --- | --- | --- |
| x4 | .000 |  |  |  |  |  |  |
| x10 | .000 | .048 |  |  |  |  |  |
| x8 | .014 | .000 | 2.735 |  |  |  |  |
| x9 | -.002 | .022 | -.331 | .531 |  |  |  |
| x6 | .000 | .000 | .081 | -.009 | .008 |  |  |
| x7 | -.001 | .079 | .104 | .113 | .013 | .254 |  |
| x11 | .000 | -.004 | .066 | .010 | .002 | -.002 | .004 |

#### Implied Correlations (Group number 1 - Default model)

|  | x4 | x10 | x8 | x9 | x6 | x7 | x11 |
| --- | --- | --- | --- | --- | --- | --- | --- |
| x4 | 1.000 |  |  |  |  |  |  |
| x10 | .000 | 1.000 |  |  |  |  |  |
| x8 | .415 | .000 | 1.000 |  |  |  |  |
| x9 | -.114 | .139 | -.275 | 1.000 |  |  |  |
| x6 | .232 | .001 | .559 | -.147 | 1.000 |  |  |
| x7 | -.057 | .710 | .125 | .308 | .293 | 1.000 |  |
| x11 | .257 | -.279 | .642 | .221 | .381 | -.074 | 1.000 |

#### Implied Means (Group number 1 - Default model)

|  | x4 | x10 | x8 | x9 | x6 | x7 | x11 |
| --- | --- | --- | --- | --- | --- | --- | --- |
|  | .089 | .430 | 4.210 | .532 | .532 | .734 | .140 |

#### Residual Covariances (Group number 1 - Default model)

|  | x4 | x10 | x8 | x9 | x6 | x7 | x11 |
| --- | --- | --- | --- | --- | --- | --- | --- |
| x4 | .000 |  |  |  |  |  |  |
| x10 | .001 | .000 |  |  |  |  |  |
| x8 | .000 | .207 | .000 |  |  |  |  |
| x9 | -.002 | -.025 | .144 | -.040 |  |  |  |
| x6 | .000 | .004 | .000 | .004 | .000 |  |  |
| x7 | .000 | .006 | .390 | -.035 | .007 | .024 |  |
| x11 | .000 | .005 | -.016 | .005 | .000 | .009 | -.001 |

#### Residual Means (Group number 1 - Default model)

|  | x4 | x10 | x8 | x9 | x6 | x7 | x11 |
| --- | --- | --- | --- | --- | --- | --- | --- |
|  | .000 | .000 | .000 | .000 | .000 | .000 | .000 |

#### Standardized Residual Covariances (Group number 1 - Default model)

|  | x4 | x10 | x8 | x9 | x6 | x7 | x11 |
| --- | --- | --- | --- | --- | --- | --- | --- |
| x4 | .000 |  |  |  |  |  |  |
| x10 | 1.129 | .000 |  |  |  |  |  |
| x8 | .000 | 4.175 | .000 |  |  |  |  |
| x9 | -1.186 | -1.137 | .844 | -.396 |  |  |  |
| x6 | -.559 | 1.644 | .005 | .482 | .005 |  |  |
| x7 | .188 | .310 | 3.415 | -.665 | 1.183 | .491 |  |
| x11 | -.774 | 2.567 | -.945 | .816 | -.095 | 2.186 | -1.001 |

#### Standardized Residual Means (Group number 1 - Default model)

|  | x4 | x10 | x8 | x9 | x6 | x7 | x11 |
| --- | --- | --- | --- | --- | --- | --- | --- |
|  | .000 | .000 | .000 | .000 | .000 | .000 | .000 |

#### Total Effects (Group number 1 - Default model)

|  | x4 | x10 | x8 | x9 | x6 | x7 |
| --- | --- | --- | --- | --- | --- | --- |
| x8 | 33.520 | .000 | .000 | .000 | .000 | .000 |
| x9 | -4.061 | .461 | -.121 | .000 | .000 | .000 |
| x6 | .997 | .000 | .030 | .001 | .000 | .000 |
| x7 | -1.400 | .001 | .055 | .002 | 1.841 | .000 |
| x11 | .778 | -.097 | .024 | .000 | .020 | .011 |

#### Standardized Total Effects (Group number 1 - Default model)

|  | x4 | x10 | x8 | x9 | x6 | x7 |
| --- | --- | --- | --- | --- | --- | --- |
| x8 | .415 | .000 | .000 | .000 | .000 | .000 |
| x9 | -.114 | .139 | -.275 | .000 | .000 | .000 |
| x6 | .232 | .001 | .559 | .007 | .000 | .000 |
| x7 | -.057 | .000 | .180 | .002 | .321 | .000 |
| x11 | .257 | -.343 | .646 | .000 | .029 | .090 |

#### Direct Effects (Group number 1 - Default model)

|  | x4 | x10 | x8 | x9 | x6 | x7 |
| --- | --- | --- | --- | --- | --- | --- |
| x8 | 33.520 | .000 | .000 | .000 | .000 | .000 |
| x9 | .000 | .461 | -.121 | .000 | .000 | .000 |
| x6 | .000 | .000 | .030 | .001 | .000 | .000 |
| x7 | -3.235 | .000 | .000 | .000 | 1.841 | .000 |
| x11 | .000 | -.097 | .024 | .000 | .000 | .011 |

#### Standardized Direct Effects (Group number 1 - Default model)

|  | x4 | x10 | x8 | x9 | x6 | x7 |
| --- | --- | --- | --- | --- | --- | --- |
| x8 | .415 | .000 | .000 | .000 | .000 | .000 |
| x9 | .000 | .139 | -.275 | .000 | .000 | .000 |
| x6 | .000 | .000 | .561 | .007 | .000 | .000 |
| x7 | -.132 | .000 | .000 | .000 | .321 | .000 |
| x11 | .000 | -.343 | .630 | .000 | .000 | .090 |

#### Indirect Effects (Group number 1 - Default model)

|  | x4 | x10 | x8 | x9 | x6 | x7 |
| --- | --- | --- | --- | --- | --- | --- |
| x8 | .000 | .000 | .000 | .000 | .000 | .000 |
| x9 | -4.061 | .000 | .000 | .000 | .000 | .000 |
| x6 | .997 | .000 | .000 | .000 | .000 | .000 |
| x7 | 1.835 | .001 | .055 | .002 | .000 | .000 |
| x11 | .778 | .000 | .001 | .000 | .020 | .000 |

#### Standardized Indirect Effects (Group number 1 - Default model)

|  | x4 | x10 | x8 | x9 | x6 | x7 |
| --- | --- | --- | --- | --- | --- | --- |
| x8 | .000 | .000 | .000 | .000 | .000 | .000 |
| x9 | -.114 | .000 | .000 | .000 | .000 | .000 |
| x6 | .232 | .001 | -.002 | .000 | .000 | .000 |
| x7 | .075 | .000 | .180 | .002 | .000 | .000 |
| x11 | .257 | .000 | .016 | .000 | .029 | .000 |

#### Modification Indices (Group number 1 - Default model)

#### Covariances: (Group number 1 - Default model)

|  |  |  | M.I. | Par Change |
| --- | --- | --- | --- | --- |

#### Variances: (Group number 1 - Default model)

|  |  |  | M.I. | Par Change |
| --- | --- | --- | --- | --- |

#### Regression Weights: (Group number 1 - Default model)

|  |  |  | M.I. | Par Change |
| --- | --- | --- | --- | --- |

#### Means: (Group number 1 - Default model)

|  |  |  | M.I. | Par Change |
| --- | --- | --- | --- | --- |

#### Intercepts: (Group number 1 - Default model)

|  |  |  | M.I. | Par Change |
| --- | --- | --- | --- | --- |

#### Minimization History (Default model)

| Iteration |  | Negative eigenvalues | Condition # | Smallest eigenvalue | Diameter | F | NTries | Ratio |
| --- | --- | --- | --- | --- | --- | --- | --- | --- |
| 0 | e | 2 |  | -.308 | 9999.000 | 273.081 | 0 | 9999.000 |
| 1 | e | 2 |  | -.393 | .549 | 139.090 | 16 | .856 |
| 2 | e | 1 |  | -.181 | .330 | 97.395 | 4 | .885 |
| 3 | e | 0 | 4444.943 |  | .400 | 70.584 | 5 | .759 |
| 4 | e | 0 | 4627.180 |  | .641 | 57.165 | 4 | .000 |
| 5 | e | 0 | 4362.578 |  | .928 | 34.572 | 1 | 1.031 |
| 6 | e | 0 | 3865.635 |  | .180 | 29.533 | 1 | 1.177 |
| 7 | e | 0 | 3639.301 |  | .043 | 28.880 | 1 | 1.119 |
| 8 | e | 0 | 3565.802 |  | .008 | 28.851 | 1 | 1.039 |
| 9 | e | 0 | 3592.048 |  | .000 | 28.851 | 1 | 1.003 |
| 10 | e | 0 | 3586.408 |  | .000 | 28.851 | 1 | 1.000 |

#### Pairwise Parameter Comparisons (Default model)

#### Variance-covariance Matrix of Estimates (Default model)

|  | par\_1 | par\_2 | par\_3 | par\_4 | par\_5 | par\_6 | par\_7 | par\_8 | par\_9 | par\_10 | par\_11 | par\_12 | par\_13 | par\_14 | par\_15 | par\_16 | par\_17 | par\_18 | par\_19 | par\_20 | par\_21 | par\_22 | par\_23 | par\_24 | par\_25 | par\_26 | par\_27 |
| --- | --- | --- | --- | --- | --- | --- | --- | --- | --- | --- | --- | --- | --- | --- | --- | --- | --- | --- | --- | --- | --- | --- | --- | --- | --- | --- | --- |
| par\_1 | .003 |  |  |  |  |  |  |  |  |  |  |  |  |  |  |  |  |  |  |  |  |  |  |  |  |  |  |
| par\_2 | .000 | 99.857 |  |  |  |  |  |  |  |  |  |  |  |  |  |  |  |  |  |  |  |  |  |  |  |  |  |
| par\_3 | .000 | .000 | .000 |  |  |  |  |  |  |  |  |  |  |  |  |  |  |  |  |  |  |  |  |  |  |  |  |
| par\_4 | .000 | .000 | .000 | .000 |  |  |  |  |  |  |  |  |  |  |  |  |  |  |  |  |  |  |  |  |  |  |  |
| par\_5 | .005 | .000 | -.001 | .000 | .205 |  |  |  |  |  |  |  |  |  |  |  |  |  |  |  |  |  |  |  |  |  |  |
| par\_6 | .012 | .000 | .003 | .000 | -.167 | 3.567 |  |  |  |  |  |  |  |  |  |  |  |  |  |  |  |  |  |  |  |  |  |
| par\_7 | .000 | .000 | .000 | .000 | .000 | .000 | .000 |  |  |  |  |  |  |  |  |  |  |  |  |  |  |  |  |  |  |  |  |
| par\_8 | .000 | .000 | .000 | .000 | .000 | .000 | .000 | .183 |  |  |  |  |  |  |  |  |  |  |  |  |  |  |  |  |  |  |  |
| par\_9 | .000 | .000 | .000 | .000 | .002 | -.005 | .000 | .007 | .001 |  |  |  |  |  |  |  |  |  |  |  |  |  |  |  |  |  |  |
| par\_10 | .000 | .000 | .000 | .000 | .000 | .000 | .000 | .002 | .000 | .000 |  |  |  |  |  |  |  |  |  |  |  |  |  |  |  |  |  |
| par\_11 | .000 | .000 | .000 | .000 | .000 | .000 | .000 | .000 | .000 | .000 | .000 |  |  |  |  |  |  |  |  |  |  |  |  |  |  |  |  |
| par\_12 | .000 | .000 | .000 | .000 | -.001 | .003 | .000 | .000 | .000 | .000 | .000 | .001 |  |  |  |  |  |  |  |  |  |  |  |  |  |  |  |
| par\_13 | .000 | .000 | .000 | .000 | .000 | .000 | .000 | .000 | .000 | .000 | .000 | .000 | .000 |  |  |  |  |  |  |  |  |  |  |  |  |  |  |
| par\_14 | .000 | .000 | .000 | .000 | .000 | .000 | .000 | .000 | .000 | .000 | .000 | .000 | .000 | .001 |  |  |  |  |  |  |  |  |  |  |  |  |  |
| par\_15 | -.001 | .000 | .000 | .000 | .000 | .000 | .000 | -.003 | .000 | .000 | .000 | .000 | .000 | .000 | .000 |  |  |  |  |  |  |  |  |  |  |  |  |
| par\_16 | .000 | -8.907 | .000 | .000 | .000 | .000 | .000 | .000 | .000 | .000 | .000 | .000 | .000 | .000 | .000 | .836 |  |  |  |  |  |  |  |  |  |  |  |
| par\_17 | -.012 | .000 | .000 | -.001 | -.020 | -.050 | .000 | -.079 | -.003 | -.001 | .000 | .000 | .000 | .000 | .004 | .000 | .094 |  |  |  |  |  |  |  |  |  |  |
| par\_18 | .000 | .000 | .000 | .000 | .000 | .000 | .000 | .000 | .000 | .000 | .000 | .000 | .000 | .000 | .000 | .000 | .000 | .000 |  |  |  |  |  |  |  |  |  |
| par\_19 | -.004 | .000 | .000 | .000 | -.094 | -.229 | .000 | .000 | -.001 | .000 | .000 | .000 | .000 | .001 | .000 | .000 | .016 | .000 | .075 |  |  |  |  |  |  |  |  |
| par\_20 | .000 | .000 | .000 | .000 | .000 | .000 | .000 | .000 | .000 | .000 | .000 | .000 | .000 | .000 | .000 | .000 | .000 | .000 | .000 | .001 |  |  |  |  |  |  |  |
| par\_21 | .000 | .000 | .000 | .000 | .000 | .000 | .000 | .000 | .000 | .000 | .000 | .000 | .000 | .000 | .000 | .000 | .000 | .000 | .000 | .000 | .000 |  |  |  |  |  |  |
| par\_22 | .000 | .000 | .000 | .000 | .000 | .000 | .000 | .000 | .000 | .000 | .000 | .000 | .000 | .000 | .000 | .000 | .000 | .000 | .000 | .000 | .000 | .190 |  |  |  |  |  |
| par\_23 | .000 | .000 | .000 | .000 | .000 | .000 | .000 | .000 | .000 | .000 | .000 | .000 | .000 | .000 | .000 | .000 | .000 | .000 | .000 | .000 | .000 | .000 | .000 |  |  |  |  |
| par\_24 | .000 | .000 | .000 | .000 | -.002 | .005 | .000 | .000 | -.001 | .000 | .000 | .001 | .000 | .000 | .000 | .000 | .000 | .000 | .001 | .000 | .000 | .000 | .000 | .008 |  |  |  |
| par\_25 | .000 | .000 | .000 | .000 | .000 | .000 | .000 | .000 | .000 | .000 | .000 | .000 | .000 | .000 | .000 | .000 | .000 | .000 | .000 | .000 | .000 | .000 | .000 | .000 | .000 |  |  |
| par\_26 | .000 | .000 | .000 | .000 | .000 | .000 | .000 | .005 | .000 | .001 | .000 | .000 | .000 | .000 | .000 | .000 | -.002 | .000 | .000 | .000 | .000 | .000 | .000 | .000 | .000 | .002 |  |
| par\_27 | .000 | .000 | .000 | .000 | .000 | .000 | .000 | .000 | .000 | .000 | .000 | .000 | .000 | .000 | .000 | .000 | .000 | .000 | .000 | .000 | .000 | .000 | .000 | .000 | .000 | .000 | .000 |

#### Correlations of Estimates (Default model)

|  | par\_1 | par\_2 | par\_3 | par\_4 | par\_5 | par\_6 | par\_7 | par\_8 | par\_9 | par\_10 | par\_11 | par\_12 | par\_13 | par\_14 | par\_15 | par\_16 | par\_17 | par\_18 | par\_19 | par\_20 | par\_21 | par\_22 | par\_23 | par\_24 | par\_25 | par\_26 | par\_27 |
| --- | --- | --- | --- | --- | --- | --- | --- | --- | --- | --- | --- | --- | --- | --- | --- | --- | --- | --- | --- | --- | --- | --- | --- | --- | --- | --- | --- |
| par\_1 | 1.000 |  |  |  |  |  |  |  |  |  |  |  |  |  |  |  |  |  |  |  |  |  |  |  |  |  |  |
| par\_2 | .000 | 1.000 |  |  |  |  |  |  |  |  |  |  |  |  |  |  |  |  |  |  |  |  |  |  |  |  |  |
| par\_3 | -.020 | .000 | 1.000 |  |  |  |  |  |  |  |  |  |  |  |  |  |  |  |  |  |  |  |  |  |  |  |  |
| par\_4 | .642 | .000 | -.169 | 1.000 |  |  |  |  |  |  |  |  |  |  |  |  |  |  |  |  |  |  |  |  |  |  |  |
| par\_5 | .190 | .000 | -.162 | .027 | 1.000 |  |  |  |  |  |  |  |  |  |  |  |  |  |  |  |  |  |  |  |  |  |  |
| par\_6 | .117 | .000 | .099 | -.017 | -.196 | 1.000 |  |  |  |  |  |  |  |  |  |  |  |  |  |  |  |  |  |  |  |  |  |
| par\_7 | .000 | .000 | .000 | .000 | .000 | .000 | 1.000 |  |  |  |  |  |  |  |  |  |  |  |  |  |  |  |  |  |  |  |  |
| par\_8 | .000 | .000 | .000 | .000 | .000 | .000 | .000 | 1.000 |  |  |  |  |  |  |  |  |  |  |  |  |  |  |  |  |  |  |  |
| par\_9 | .014 | .000 | -.697 | .118 | .113 | -.069 | .000 | .439 | 1.000 |  |  |  |  |  |  |  |  |  |  |  |  |  |  |  |  |  |  |
| par\_10 | .000 | .000 | .000 | .000 | .000 | .000 | .000 | .215 | .000 | 1.000 |  |  |  |  |  |  |  |  |  |  |  |  |  |  |  |  |  |
| par\_11 | .000 | .000 | -.002 | .000 | .002 | .000 | .000 | .000 | .002 | .000 | 1.000 |  |  |  |  |  |  |  |  |  |  |  |  |  |  |  |  |
| par\_12 | -.011 | .000 | .511 | -.086 | -.086 | .051 | .000 | .000 | -.356 | .000 | -.001 | 1.000 |  |  |  |  |  |  |  |  |  |  |  |  |  |  |  |
| par\_13 | .000 | .000 | .000 | .000 | .000 | .000 | .275 | .000 | .000 | .000 | .000 | .000 | 1.000 |  |  |  |  |  |  |  |  |  |  |  |  |  |  |
| par\_14 | .000 | .000 | .000 | .000 | .000 | .000 | .000 | .000 | .000 | .000 | .000 | .000 | .000 | 1.000 |  |  |  |  |  |  |  |  |  |  |  |  |  |
| par\_15 | -.490 | .000 | .101 | -.760 | -.016 | .010 | .000 | -.354 | -.485 | .000 | .000 | .051 | .000 | .000 | 1.000 |  |  |  |  |  |  |  |  |  |  |  |  |
| par\_16 | .000 | -.975 | .000 | .000 | .000 | .000 | .000 | .000 | .000 | .000 | .000 | .000 | .000 | .000 | .000 | 1.000 |  |  |  |  |  |  |  |  |  |  |  |
| par\_17 | -.738 | .000 | .015 | -.474 | -.141 | -.086 | .000 | -.601 | -.274 | -.129 | .000 | .008 | .000 | .000 | .630 | .000 | 1.000 |  |  |  |  |  |  |  |  |  |  |
| par\_18 | .000 | .000 | .000 | .000 | .000 | .000 | .000 | .000 | .000 | .000 | .000 | .000 | .000 | .000 | .000 | .000 | .000 | 1.000 |  |  |  |  |  |  |  |  |  |
| par\_19 | -.240 | .000 | .082 | -.014 | -.760 | -.444 | .000 | .000 | -.057 | .000 | -.001 | .045 | .000 | .178 | .008 | .000 | .197 | .000 | 1.000 |  |  |  |  |  |  |  |  |
| par\_20 | .000 | .000 | .000 | .000 | .000 | .000 | -.918 | .000 | .000 | .000 | .000 | .000 | -.478 | .000 | .000 | .000 | .000 | .000 | .000 | 1.000 |  |  |  |  |  |  |  |
| par\_21 | .000 | .000 | .000 | .000 | .000 | .000 | .000 | .000 | .000 | .000 | .000 | .000 | .000 | .000 | .000 | .000 | .000 | .000 | .000 | .000 | 1.000 |  |  |  |  |  |  |
| par\_22 | .000 | .000 | .000 | .000 | .000 | .000 | .000 | .000 | .000 | .000 | .000 | .000 | .000 | .000 | .000 | .000 | .000 | .000 | .000 | .000 | .000 | 1.000 |  |  |  |  |  |
| par\_23 | .000 | .000 | .000 | .000 | .000 | .000 | .000 | .000 | .000 | .848 | .000 | .000 | .000 | .000 | .000 | .000 | .000 | .000 | .000 | .000 | .000 | .000 | 1.000 |  |  |  |  |
| par\_24 | -.006 | .000 | .306 | -.052 | -.048 | .030 | .000 | .000 | -.214 | .000 | .667 | .509 | .000 | .000 | .031 | .000 | .004 | .000 | .024 | .000 | .000 | .000 | .000 | 1.000 |  |  |  |
| par\_25 | .000 | .000 | .000 | .000 | .000 | .000 | .000 | .000 | .000 | .000 | .000 | .000 | .000 | .000 | .000 | .000 | .000 | .000 | .000 | .000 | .000 | .000 | .000 | .000 | 1.000 |  |  |
| par\_26 | .000 | .000 | .000 | .000 | -.002 | .000 | .000 | .284 | .000 | .848 | .000 | .244 | .000 | .000 | .000 | .000 | -.170 | .000 | .002 | .000 | .000 | .000 | .560 | .073 | .000 | 1.000 |  |
| par\_27 | .000 | .000 | -.003 | .000 | .000 | .000 | .000 | .000 | .002 | .000 | .786 | -.001 | .000 | .000 | .000 | .000 | .000 | .000 | .000 | .000 | .000 | .000 | .000 | .380 | .000 | .000 | 1.000 |

#### Model Fit Summary

#### CMIN

| Model | NPAR | CMIN | DF | P | CMIN/DF |
| --- | --- | --- | --- | --- | --- |
| Default model | 27 | 28.851 | 8 | .000 | 3.606 |
| Saturated model | 35 | .000 | 0 |  |  |
| Independence model | 14 | 182.478 | 21 | .000 | 8.689 |

#### Baseline Comparisons

| Model | NFI Delta1 | RFI rho1 | IFI Delta2 | TLI rho2 | CFI |
| --- | --- | --- | --- | --- | --- |
| Default model | .842 | .585 | .880 | .661 | .871 |
| Saturated model | 1.000 |  | 1.000 |  | 1.000 |
| Independence model | .000 | .000 | .000 | .000 | .000 |

#### Parsimony-Adjusted Measures

| Model | PRATIO | PNFI | PCFI |
| --- | --- | --- | --- |
| Default model | .381 | .321 | .332 |
| Saturated model | .000 | .000 | .000 |
| Independence model | 1.000 | .000 | .000 |

#### NCP

| Model | NCP | LO 90 | HI 90 |
| --- | --- | --- | --- |
| Default model | 20.851 | 8.085 | 41.187 |
| Saturated model | .000 | .000 | .000 |
| Independence model | 161.478 | 122.012 | 208.418 |

#### FMIN

| Model | FMIN | F0 | LO 90 | HI 90 |
| --- | --- | --- | --- | --- |
| Default model | .534 | .386 | .150 | .763 |
| Saturated model | .000 | .000 | .000 | .000 |
| Independence model | 3.379 | 2.990 | 2.259 | 3.860 |

#### RMSEA

| Model | RMSEA | LO 90 | HI 90 | PCLOSE |
| --- | --- | --- | --- | --- |
| Default model | .220 | .137 | .309 | .001 |
| Independence model | .377 | .328 | .429 | .000 |

#### AIC

| Model | AIC | BCC | BIC | CAIC |
| --- | --- | --- | --- | --- |
| Default model | 82.851 | 92.242 |  |  |
| Saturated model | 70.000 | 82.174 |  |  |
| Independence model | 210.478 | 215.347 |  |  |

#### ECVI

| Model | ECVI | LO 90 | HI 90 | MECVI |
| --- | --- | --- | --- | --- |
| Default model | 1.534 | 1.298 | 1.911 | 1.708 |
| Saturated model | 1.296 | 1.296 | 1.296 | 1.522 |
| Independence model | 3.898 | 3.167 | 4.767 | 3.988 |

#### HOELTER

| Model | HOELTER .05 | HOELTER .01 |
| --- | --- | --- |
| Default model | 30 | 38 |
| Independence model | 10 | 12 |

#### Execution time summary

|  |  |
| --- | --- |
| Minimization: | .011 |
| Miscellaneous: | .065 |
| Bootstrap: | .000 |
| Total: | .076 |
