## supporting information for "The past and potential antagonistic effects of fishing ban against nutrient loading control on eutrophication restoration in Lake Hongze" (from another version of this preprint: version 5, 2026-09-09)

**Table of Content**

**Figures:**

Figure S1. PCLake model structure (lake part).

Figure S2. Monthly observations of water quality.

Figure S3. Zoobenthos composition in biomass and total dry weight of three zones in Lake Hongze in the period of 2016-2020.

Figure S4. The simulation using original model before calibration (line) and in-lake observations of different zones in Lake Hongze between January 2016 and December 2019 (dots).

**Tables:**

Table S1. Average river inflow of Lake Hongze in the period of January 2016 and December 2019.

Table S2. The performance of original and calibrated model for Lake Hongze, as well as the model performances of other studies.

Table S3. Standardized path coefficients from SEM path analysis between lake components.

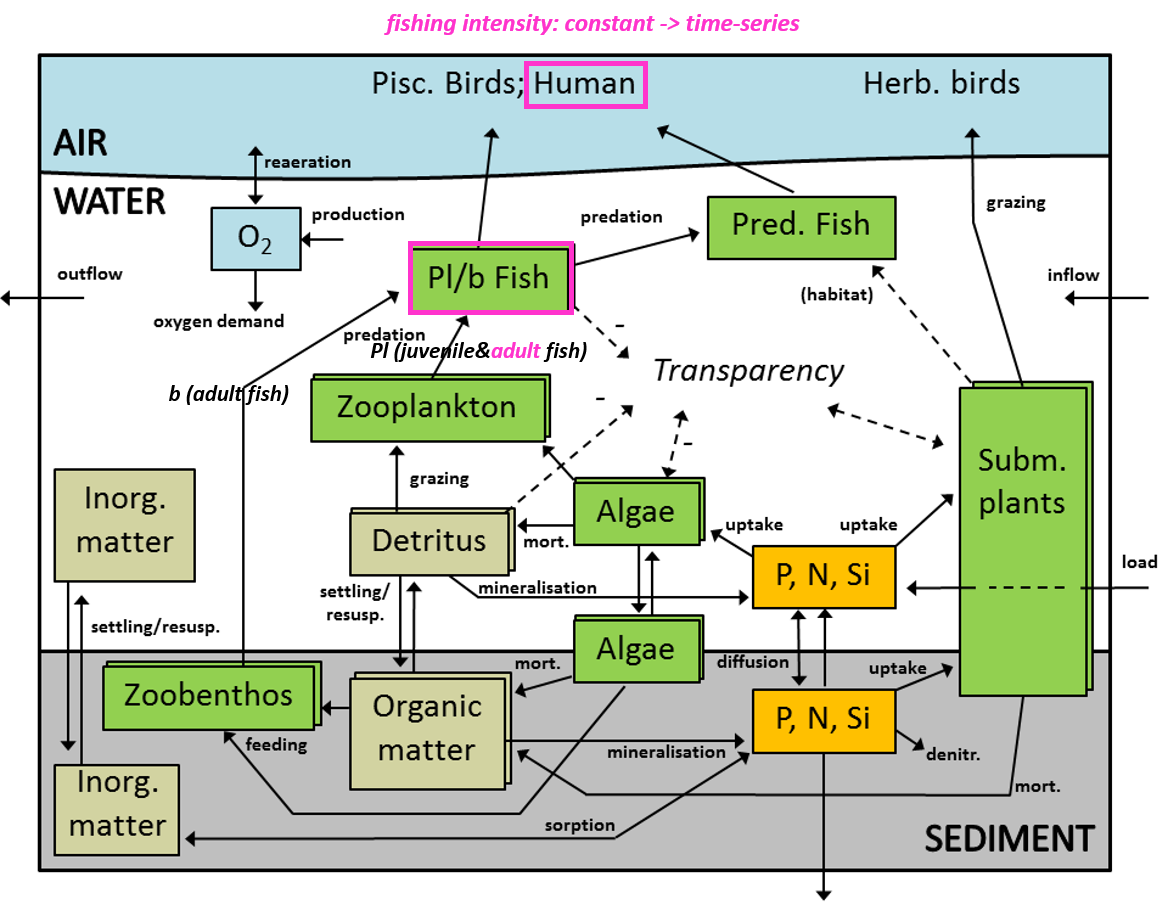

**Figure S1. PCLake model structure (lake part). Cyanobacteria, diatoms and other algae are three distinguished functional groups of phytoplankton. Fish is split into a juvenile (zooplanktivorous) and an adult (benthivorous) subgroup (Janse, 2005). The pink-colored text and box denote the two modifications implemented in the present study, namely: (i) the incorporation of zooplanktivorous adult whitefish, and (ii) the redefinition of fishing intensity from a constant first-order rate to time-series forcing data.**

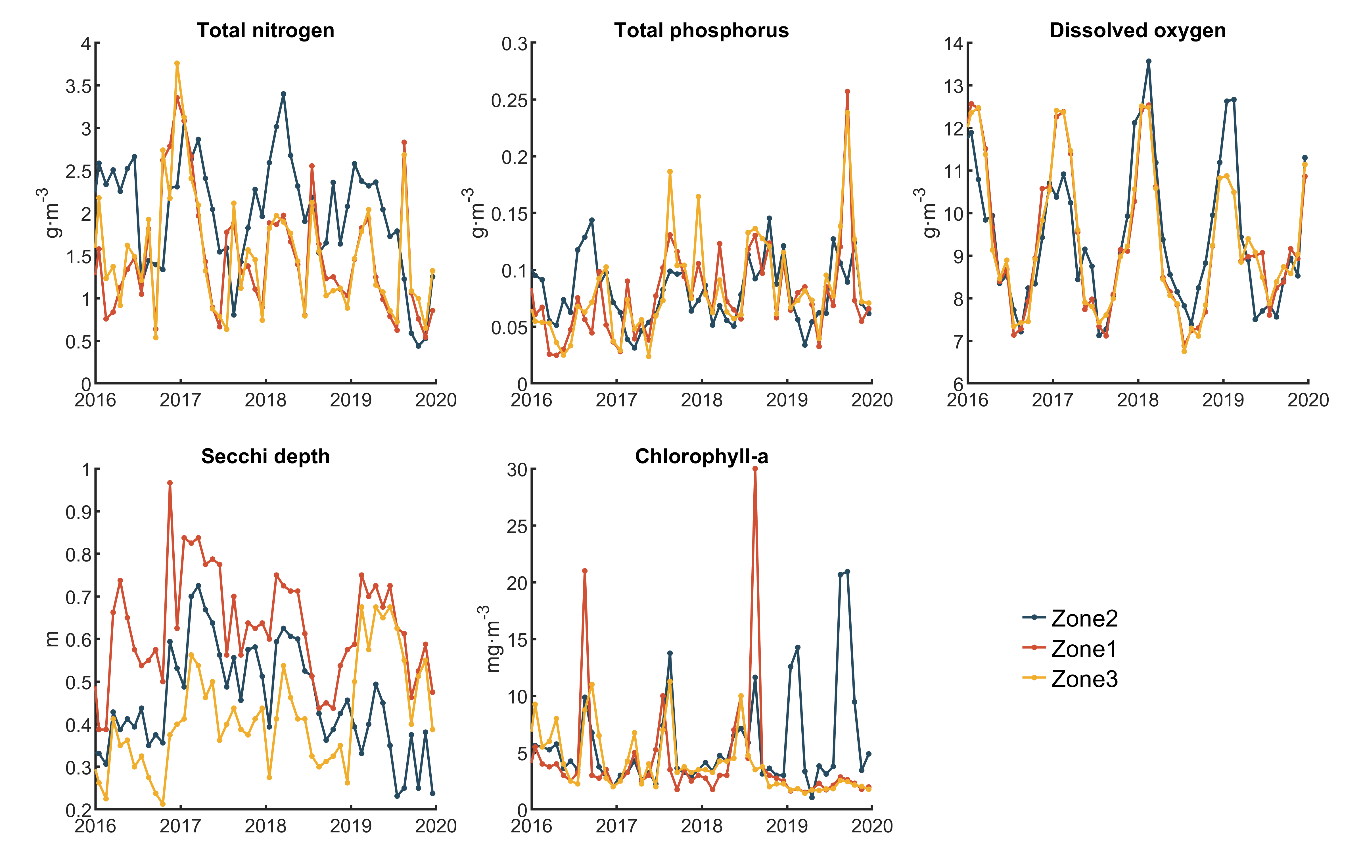

**Figure S2. Monthly observations of water quality. Zone 1 (Chengzi District) is in the north and Zone 3 (Lihewa District) in the west, and Zone 2 is the open water area.**

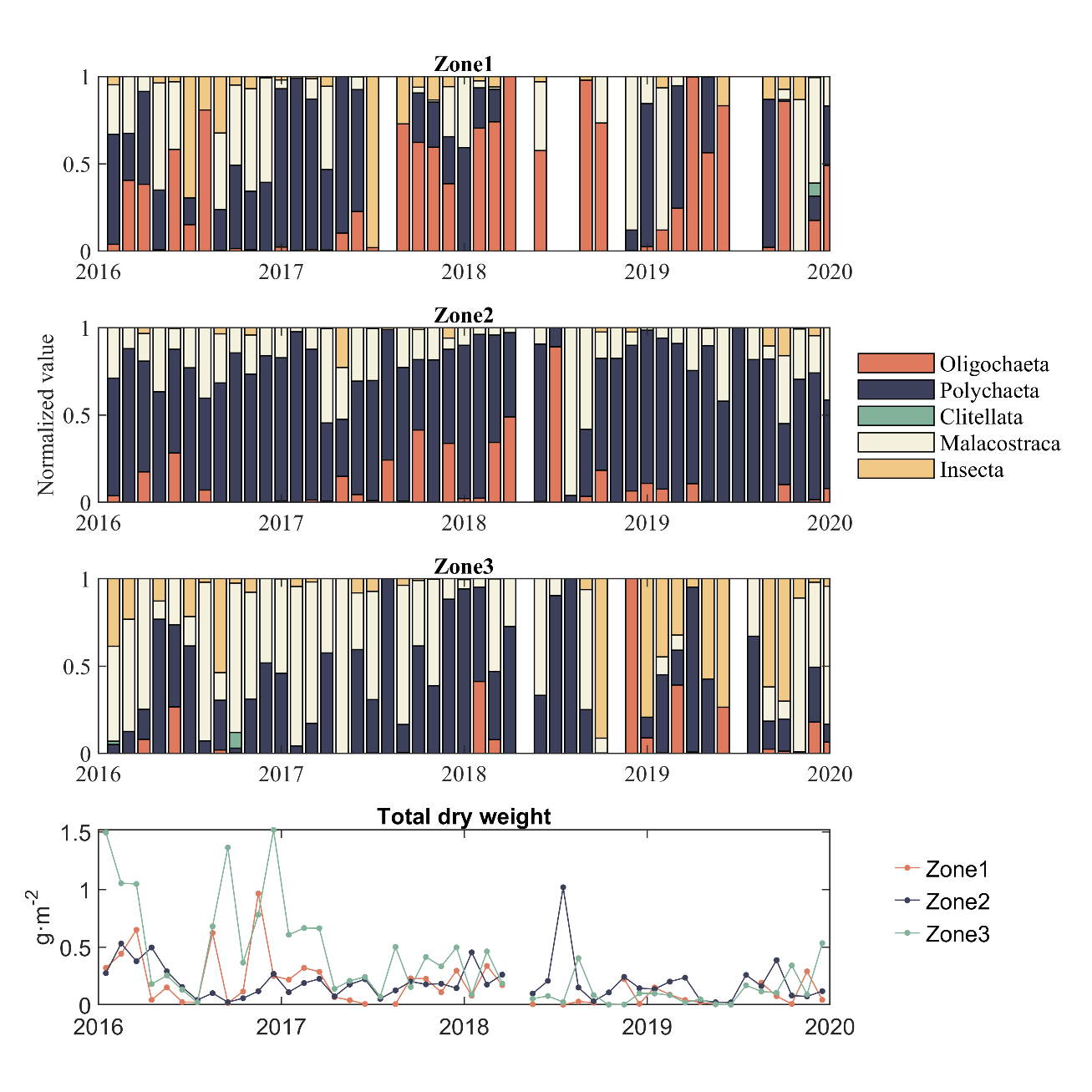

**Figure S3. Zoobenthos composition in biomass and total dry weight of three zones in Lake Hongze in the period of 2016-2020.**

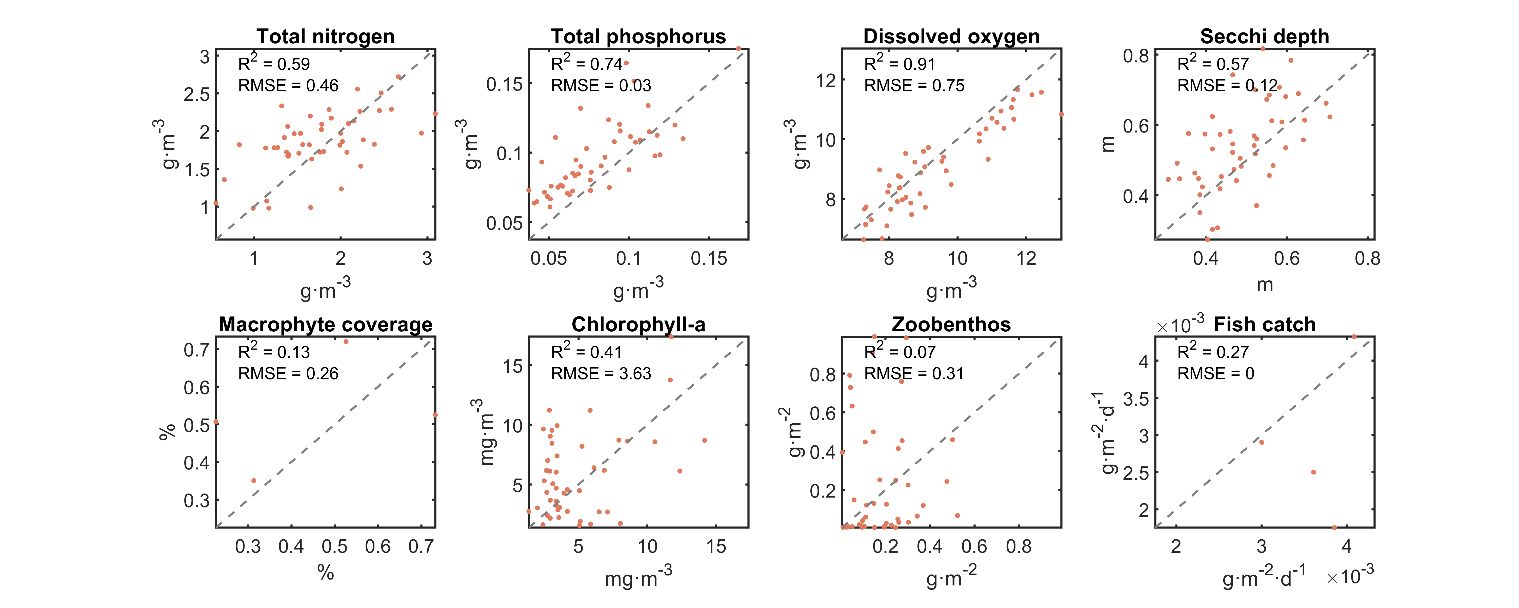

**Figure S4 - The result of PCLake model calibration between 2016 and 2020. Monthly observational data from Lake Hongze comprised total nitrogen, total phosphorus, dissolved oxygen, Secchi depth, chlorophyll‑a, and zoobenthos, while macrophyte coverage and fish harvest were recorded annually. Data from different lake zones are presented; however, the coefficient of determination (R2) and root mean square error (RMSE) were calculated based on whole-lake averages and corresponding model simulations.**

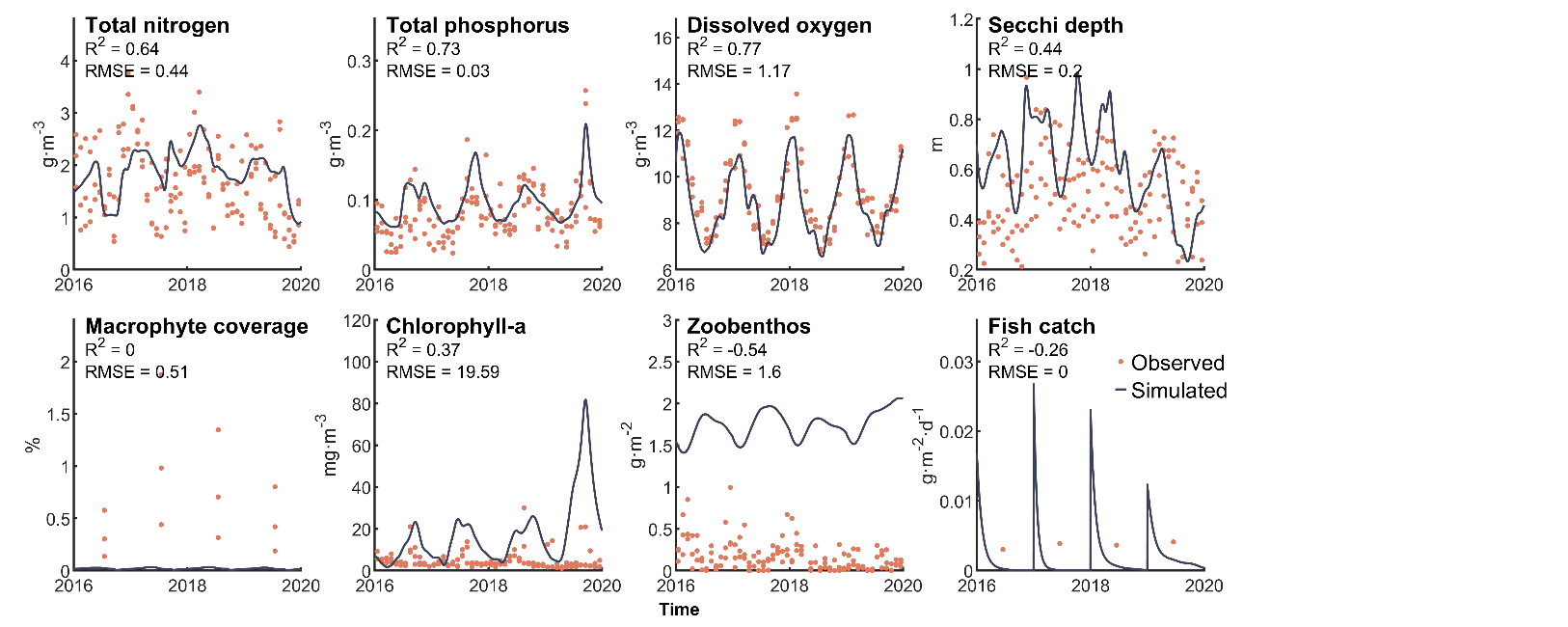

**Figure S5. The simulation using original model before calibration (line) and in-lake observations of different zones in Lake Hongze between January 2016 and December 2019 (dots). The coefficient of determination (R2) and root mean squared error (RMSE) were calculated according to observations of whole-lake average and model simulations.**

**Table S****1. Average river inflow of Lake Hongze in the period of January 2016 and December 2019.**

| Month | Chihe | Huaihongxinhe* | Huaihongxinhe** | Huaihe | Xinbianhe | Suihe | Laosuihe |
| --- | --- | --- | --- | --- | --- | --- | --- |
| 1 | 1.08 | 0.89 | 26.47 | 610.75 | 0.36 | 11.81 | 0.00 |
| 2 | 0.33 | 2.01 | 8.73 | 526.75 | 0.20 | 7.46 | 0.00 |
| 3 | 0.73 | 1.70 | 19.05 | 618.25 | 0.00 | 6.88 | 0.04 |
| 4 | 0.00 | 3.70 | 9.15 | 481.75 | 0.08 | 4.23 | 0.00 |
| 5 | 24.70 | 0.79 | 18.19 | 830.50 | 0.00 | 2.50 | 0.19 |
| 6 | 16.75 | 14.74 | 49.66 | 911.00 | 0.24 | 10.02 | 3.75 |
| 7 | 24.08 | 37.51 | 85.88 | 1408.75 | 0.13 | 25.19 | 7.00 |
| 8 | 15.28 | 116.10 | 132.75 | 864.03 | 67.25 | 91.48 | 12.73 |
| 9 | 13.38 | 17.63 | 78.12 | 807.75 | 7.89 | 13.50 | 0.99 |
| 10 | 19.93 | 43.40 | 109.63 | 1380.18 | 4.92 | 41.13 | 3.05 |
| 11 | 9.68 | 3.83 | 37.95 | 1076.10 | 0.42 | 12.60 | 0.00 |
| 12 | 1.14 | 0.00 | 15.81 | 500.25 | 0.00 | 10.26 | 0.00 |
| Total | 127.06 | 242.28 | 591.38 | 10016.05 | 81.49 | 237.03 | 27.74 |
| % | 1.1 | 2.1 | 5.2 | 88.5 | 0.7 | 2.1 | 0.2 |

* Shuanggou station ** Xiacaowan station. The data unit for inflow is m^3^∙s^-1^.

**Table S2 - Composition of dominant fish species in Lake Hongze in 2018.**

| Feeding habits | Fish Species | N% | W% | IRI | Total (N%) |
| --- | --- | --- | --- | --- | --- |
| zooplanktivorous | *Aristichthys nobilis* | 2.0 | 41.1 | 4315.7 | 27.6 |
|  | *Coilia nasus* | 25.6 | 4.4 | 3002.8 |  |
| omnivores | *Hemiculter leucisculus* | 27.3 | 7.3 | 3462.2 | 45.9 |
|  | *Carassius auratus* | 16.7 | 17.7 | 3447.2 |  |
|  | *Pseudorasbora parva* | 1.9 | 0.1 | 200.8 |  |
| predatory | *Pelteobagrus nitidus* | 2.1 | 0.5 | 265.1 | 21.6 |
|  | *Pelteobagrus fulvidraco* | 8.5 | 4.3 | 1278.5 |  |
|  | *Cultrichthys erythropterus* | 11 | 9.7 | 2063.8 |  |
| herbivores | *Ctenopharyngodon idellus* | 0.1 | 2.8 | 277.3 | 0.4 |
|  | *Hypophthalmichthys molitrix* | 0.3 | 4.9 | 523 |  |

Note: N%, W% and IRI are the percentage of numbers, weight and index of relative importance of fish species. Data was obtained from previous study (Mao et al. 2019) and has been reclassified by feeding habits.

**Table S3. The performance of original and calibrated model for Lake Hongze, as well as the model performances of other studies.**

|  |  | **R2** |  |  | **RMSE** |  |
| --- | --- | --- | --- | --- | --- | --- |
|  | Original | Modified | Other studies* | Original | Modified | Other studies** |
| Total nitrogen | 0.64 | 0.65 | 0.03-0.89 | 0.44 | 0.43 | 0.81-1.64 |
| Total phosphorus | 0.73 | 0.72 | 0-0.63 | 0.03 | 0.03 | 0.02-0.16 |
| Dissolved oxygen | 0.77 | 0.80 | 0-0.94 | 1.17 | 1.03 | 0.11-2.24 |
| Secchi depth | 0.44 | 0.40 | 0.18-0.63 | 0.20 | 0.13 | 0.12-0.25 |
| Macrophyte coverage | 0 | 0.49 | 0.05-0.94 | 0.51 | 0.19 | 0.35-48.1 |
| Chlorophyll-a | 0.37 | 0.39 | 0.34 | 19.59 | 3.72 | 4.95 |
| Zoobenthos | -0.54 | 0.27 | 0-0.96 | 1.6 | 0.3 |  |
| Fish catch | -0.26 | 0.78 | 0.18-0.83 | 0 | 0 |  |

* *Number from other studies were collected by Qin et al. (2025)*

Qin, B., Xu, M., Zhu, K., et al. 2025. Climate warming will significantly affect future restoration and level of ecosystem services in Lake Erhai. ECOLOGICAL MODELLING, 503, 111067. https://doi.org/https://doi.org/10.1016/j.ecolmodel.2025.111067

*** Number was collected in this study from below literatures:*

Fragoso, C. R., Marques, D. M. L. M., Ferreira, T. F., et al. 2011. Potential effects of climate change and eutrophication on a large subtropical shallow lake. ENVIRONMENTAL MODELLING & SOFTWARE, 26(11), 1337-1348. https://doi.org/10.1016/j.envsoft.2011.05.004

Kong, X. Z., Dong, L., He, W., et al. 2015. Estimation of the long-term nutrient budget and thresholds of regime shift for a large shallow lake in China. ECOLOGICAL INDICATORS, 52, 231-244. https://doi.org/10.1016/j.ecolind.2014.12.005

Kong, X., Zhan, Q., Boehrer, B., et al. 2019. High frequency data provide new insights into evaluating and modeling nitrogen retention in reservoirs. WATER RESEARCH, 166. https://doi.org/10.1016/j.watres.2019.115017

Andersen, T. K., Nielsen, A., Jeppesen, E., et al. 2020. Predicting ecosystem state changes in shallow lakes using an aquatic ecosystem model: Lake Hinge, Denmark, an example. ECOLOGICAL APPLICATIONS, 30(7). https://doi.org/10.1002/eap.2160

Qin, B., Kong, X., Wang, R., et al. 2022. Lake restoration time of Lake Taibai (China): a case study based on paleolimnology and ecosystem modeling. J Paleolimnol, 68, 25–38. https://doi.org/https://doi.org/10.1007/s10933-020-00165-7

Shi, T., Chen, Y., Zhang, H., et al. 2023. Numerical study on regime shifts in an urban subtropical shallow lake: Xinglong Lake, China. ECOLOGICAL INDICATORS, 154. https://doi.org/10.1016/j.ecolind.2023.110600

Zhang, X., Yi, Y., Cao, Y., et al. 2023. Disentangling the effects of phosphorus loading on food web stability in a large shallow lake. JOURNAL OF ENVIRONMENTAL MANAGEMENT, 328. https://doi.org/10.1016/j.jenvman.2022.116991
