## Supplementary material for "The past and potential antagonistic effects of fishing ban against nutrient loading control on eutrophication restoration in Lake Hongze": author response (from another version of this preprint: version 5, 2026-09-09)

Summary: We thank the reviewers for their constructive comments. In response to the concern regarding SEM p-values, we have re-analyzed the model by including missing links identified via modification indices. The updated SEM shows improved fit, while our core conclusions remain unchanged. Revisions have been made in Section 3.2 (lines 328–338 and Figure 3B) and the Discussion (lines 478–479). We have also corrected minor table reference errors (lines 307, 463, and 468). We believe the manuscript is now suitable for publication.

Review by Benoit Pichon, 02 Aug 2026 14:49

I thank the authors for their constructive responses, which really increased the quality of their manuscript. The text reads better and its easier to follow the story: well done !

I would have one remaining comment regarding the statistical analyses. The authors provide the p-value associated with their SEM. However, if I'm not wrong, these p-values suggest that they are one (or multiple) links that are significant but not included in the current model (such that p-value is below 0.05). Could the authors clarify that, since this can strongly affect the other relationships evaluated in the SEM? This might need to include some missing links or correlation structure in the current SEM in Fig. 3.

Response: We sincerely thank you for your positive and constructive feedback. We fully agree with your comment regarding the p-values associated with our SEM. We acknowledge that this issue arose partly due to data standardization and the omission of certain ecologically plausible pathways. To address this, we have re‑analyzed the SEM by incorporating modification indices to identify and include missing links that are both statistically justified and ecologically meaningful. The updated models now show improved fit, while the core conclusions regarding the antagonistic effects between fishing bans and nutrient control remain unchanged. Accordingly, we have revised the following sections of the manuscript:

Section 3.2 (lines 328–338 and Figure 3B): The SEM results have been updated with the new path coefficients, fit statistics, and a brief explanation of the added links.

Discussion (lines 478–479): We have clarified the model revision process and noted that the improved fit supports the robustness of our main findings.

We are confident that these revisions adequately address your concern and further strengthen the validity of our SEM analysis. We hope that the updated version meets your expectations.

Other modification:

Response: In addition, we have corrected several minor errors in table references throughout the manuscript (lines 307, 463, and 468).
